# Molecular Determinants of Allosteric Inhibitor Affinity and Selectivity in PDE5

**DOI:** 10.64898/2026.03.16.712116

**Authors:** Jyoti Verma, Harish Vashisth

## Abstract

Allosteric inhibitors targeting unique regulatory pockets provide an opportunity for selective modulation of proteins within the phosphodiesterase (PDE) family. In this work, we investigate the molecular determinants for binding of an evodiamine derivative (EVO) to PDE5 and the structural basis for differential recognition of this inhibitor in PDE6 isoforms. We integrated structural modeling, all-atom molecular dynamics (MD) simulations, and extensive alchemical free-energy calculations to quantify residue-level contributions to inhibitor binding in PDE5 through systematic alanine scanning and PDE6-derived mutations. Thermodynamic analysis based on free energy identified key residues that govern EVO binding through a network of hydrogen bonds and hydrophobic contacts. Specifically, we examined structural changes associated with allosteric pocket variations in PDE5, identified key structural elements, and evaluated the role of chemical substituents of EVO in allosteric inhibition. Moreover, PDE6-derived substitutions revealed isoform specific effects, with PDE6-rod substitutions resulting in stronger destabilization than PDE6-cone substitutions. Together, these results define the energetic and structural basis for allosteric inhibition of PDE5 and provide mechanistic insights for the rational design of selective PDE inhibitors.

**TOC Graphic:** 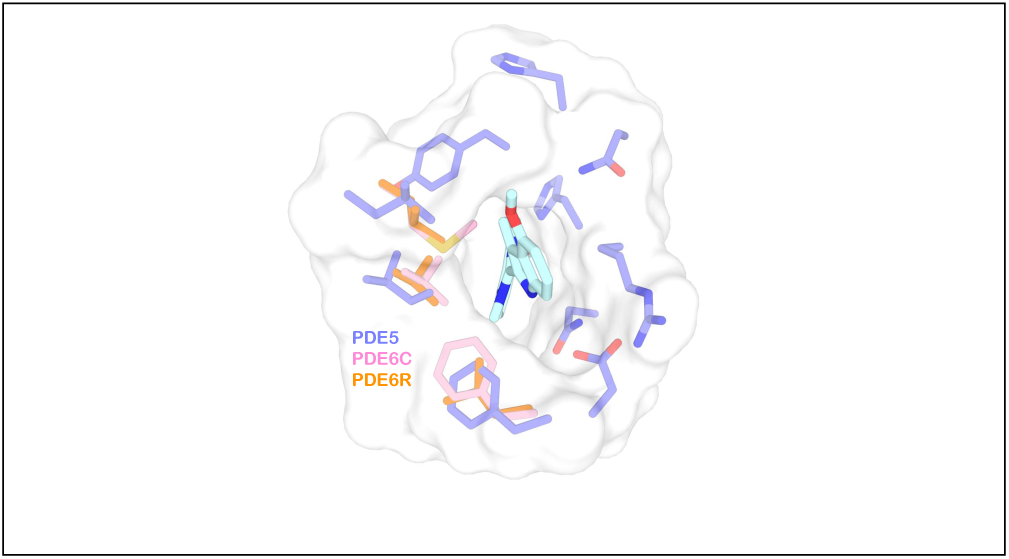

## Introduction

Phosphodiesterases (PDEs) are a superfamily of phosphohydrolases that hydrolyze the second messengers adenosine and guanosine 3*^′^,* 5*^′^*-cyclic monophosphates (cAMP and cGMP) into their inactive 5*^′^*-monophosphate forms.^1–3^ By regulating intracellular concentrations of cAMP and cGMP, PDEs act as key modulators of diverse signaling cascades underlying physiological processes including smooth muscle contraction, inflammation, memory, vision, and immune responses.^4^ The mammalian genome encodes 11 PDE families (PDE1–PDE11), each with multiple isoforms, differing in tissue distribution, substrate selectivity, regulatory mechanisms, and inhibitor sensitivity. ^5^ PDE families are broadly classified according to their substrate specificity:^6^ (i) cAMP specific (PDE4, PDE7, PDE8),^7,8^ (ii) cGMP specific (PDE5, PDE6, PDE9),^9–11^ and (iii) dual-substrate which can hydrolyze both cAMP and cGMP (PDE1, PDE2, PDE3, PDE10, PDE11).^7,12,13^ This classification highlights how different PDE isoforms have evolved to regulate distinct signaling pathways. PDEs have also been validated as drug targets, with inhibitors already in clinical use for erectile dysfunction, chronic obstructive pulmonary disease (COPD), psoriasis, and heart failure. ^14–16^ However, understanding the structural determinants of substrate recognition and allosteric regulation in PDEs is critical for designing the next-generation of selective modulators with improved therapeutic efficacy and reduced side effects.

Among PDEs, PDE5 and PDE6 are two closely related but functionally divergent enzymes. PDE5 is a cGMP specific enzyme that plays a central role in smooth muscle relaxation and vascular signaling. ^17^ Dysregulation of PDE5 is implicated in cardiovascular diseases, pulmonary hypertension, and erectile dysfunction, making it a clinically important therapeutic target.^18–21^ PDE6 is also a cGMP-specific enzyme, but unlike PDE5, it is primarily expressed in the retina, where it plays a critical role in the visual transduction cascade. ^22,23^ Despite their distinct physiological functions, both enzymes are highly selective for cGMP and share structural similarities, leading to potential inhibitor cross-reactivity with implications for drug selectivity and off-target effects. Various PDE5 inhibitors (e.g., sildenafil) show cross-reactivity with PDE6, leading to visual side effects such as altered color perception or increased light sensitivity in patients undergoing treatment for erectile dysfunction or pulmonary hypertension. ^24,25^ This off-target inhibition highlights the challenge of achieving selectivity among closely related PDEs. ^26^ Therefore, it is crucial to understand subtle structural and dynamical differences between PDE5 and PDE6 at the molecular level.^27^ Current therapeutic strategies against PDE5 are competitive inhibitors (orthosteric inhibitors) that bind to the catalytic site of PDE5, including sildenafil, tadalafil, and vardenafil.^28–30^ However, these inhibitors are often associated with off-target effects. In contrast, allosteric modulation of PDE5 has emerged as an alternative strategy to overcome these challenges.^31,32^ Unlike orthosteric inhibitors, allosteric inhibitors bind to regulatory sites distinct from the catalytic pocket, thereby indirectly modulating the enzyme activity. In PDE5, allosteric pockets within regulatory GAF domains have been extensively characterized and shown to influence conformational dynamics and catalytic efficiency.^33^ In contrast, the catalytic domain, a primary target of orthosteric inhibitors, has not been targeted by allosteric modulators. However, the discovery of evodiamine derivatives as selective allosteric inhibitors of PDE5 illustrates the potential to target allosteric sites within the catalytic domain.^32^

In particular, the structure of PDE5 cocrystalized with an evodiamine derivative, (S)-7e, revealed a novel allosteric pocket (Figure 1).^32^ Binding of this evodiamine derivative (hereafter termed EVO) to the allosteric pocket triggered large conformational rearrangements, including a ∼24 Å displacement of the H-loop, effectively occluding the catalytic site and inhibiting enzyme activity (Figure S1). The EVO-based allosteric inhibitor series showed 570-fold selectivity towards PDE5 over cone-PDE6 (PDE6C).^32^ Furthermore, these allosteric inhibitors also showed *in vivo* efficacy in preclinical models of pulmonary hypertension, supporting their relevance as leads for selective modulation of PDE5. ^32^ Despite these promising experimental findings, the energetic contributions of residues involved in binding of EVO and hence the basis of its selectivity for PDE5 is yet to be fully resolved. It is also unclear how individual residues within the allosteric pocket differentially contribute to binding affinity and how sequence variations in homologous PDE6 isoforms (cone PDE6; PDE6C, and rod PDE6; PDE6R) influence inhibitor recognition.

**Figure 1:**
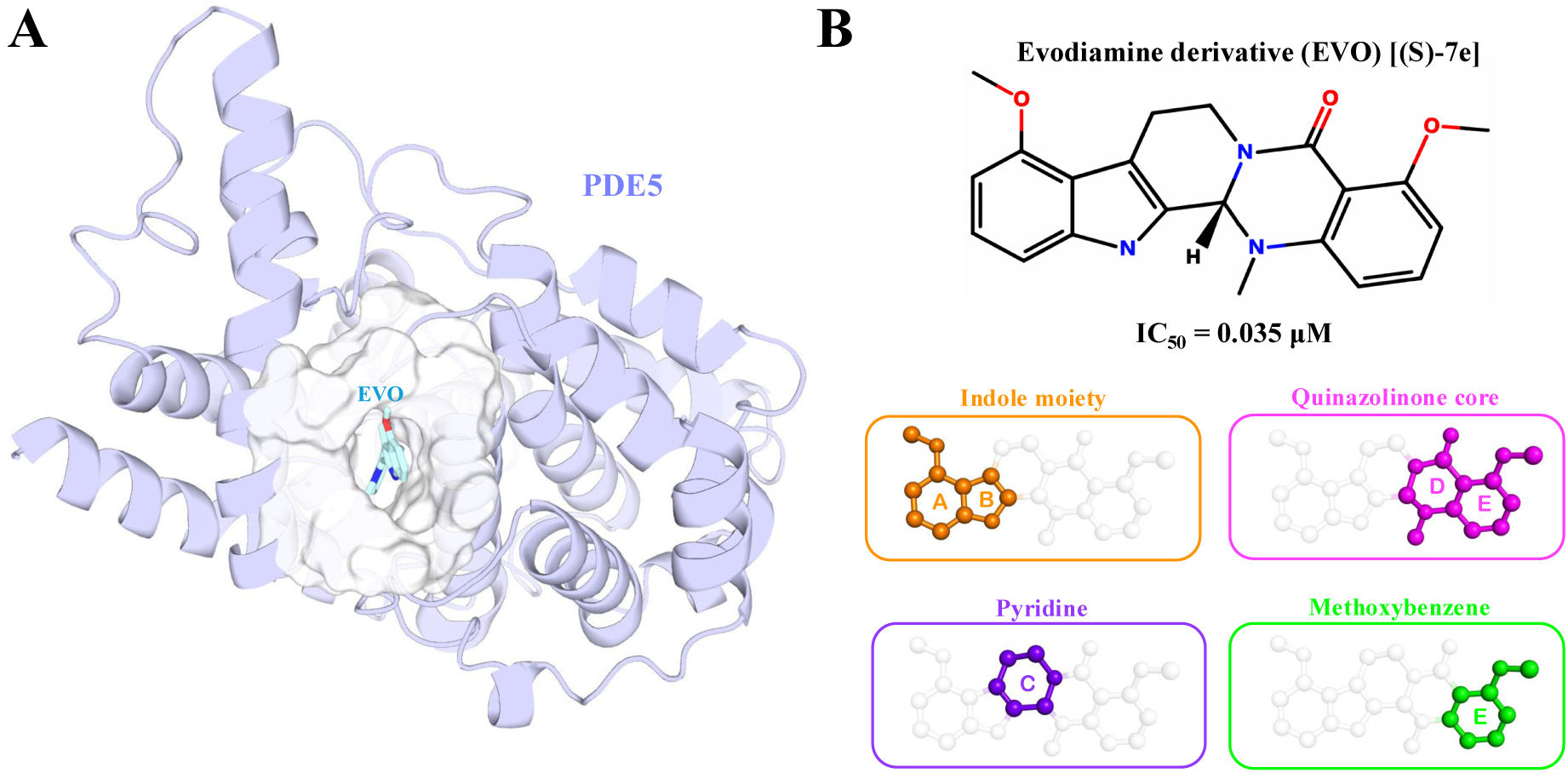
(A) Structure of PDE5 in complex with an evodiamine derivative, S-7e (termed as EVO in this study); PDB ID 6VBI. The structure shown here is the modeled energy minimized structure of the complex. (B) The chemical structure of EVO and its reported experimental activity (IC_50_) toward PDE5. The lower panels show a structural classification of chemical moieties in EVO, highlighted in different colors. Ring A and B: Indole moiety (orange); Ring C: Fused pyridine ring (purple); Ring D and E: Quinazolinone core (magenta); Ring E: Fused methoxybenzene (of Quinazolinone core) (green).

To understand the molecular basis of allosteric inhibition and selectivity, it is therefore critical to evaluate the energetic contributions of residues in PDE6 isoforms that correspond to PDE5 residues involved in binding of the allosteric inhibitor (EVO). In this work, we investigate the molecular basis of allosteric inhibition in PDE5 and provide insights into energetics of EVO binding. Specifically, we performed alchemical free energy calculations on key allosteric pocket residues in PDE5 to identify those residues that are critical to inhibitor binding. Additionally, residues that differed from PDE5 in PDE6 isoforms were mutated to replicate a PDE6-like environment, allowing us to assess the structural and energetic determinants of selectivity.

## Materials and Methods

### System Setup: Structural Modeling of PDE5 and Simulation Details

The atomic coordinates of PDE5 bound to the allosteric inhibitor EVO were obtained from the protein data bank (PDB ID:6VBI).^32^ The residues 789-809, located within the *α*14-helix, were unresolved in the crystal structure of PDE5-EVO complex (Figure S1). Based on the reference sequence (Uniprot ID:O76074), we first obtained a complete structural model of PDE5 using the AlphaFold3 server.^34,35^ The AlphaFold predicted full-length PDE5 model showed high confidence for the missing region (784-809) with a predicted TM-score (pTM) of 0.95. Then, we extracted the atomic coordinates of the missing region (784-809) from the AlphaFold-predicted structure and integrated it with the crystal structure of PDE5 using the *psfgen* tool in VMD.^36^

After building a complete structural model of PDE5, we performed energy minimization on the modeled structure to ensure proper geometry of the backbone dihedral angles and the peptide-bond lengths. The complete PDE5 structure in complex with the allosteric inhibitor was solvated with explicit TIP3P water molecules in a periodic simulation domain. ^37^ The system was neutralized and adjusted to a physiological ionic strength of 150 mM by adding Na^+^ and Cl*^−^* ions to the bulk solvent. Finally, the solvated and ionized structure was minimized for 10000 steps using the conjugate gradient algorithm in NAMD.^38^ For energy minimization, we fixed the backbone atoms of the protein except for residues 780–813 comprising the modeled region (784-809) and four adjacent residues on each side, which were kept flexible to facilitate structural relaxation. The inclusion of adjacent residues was intended to ensure smooth backbone continuity at the junction between the modeled and experimentally resolved regions. All non-hydrogen atoms of the inhibitor were also fixed to retain their positions as observed in the crystal structure. The minimized structure was assessed by analyzing the Ramachandran plot outliers,^39^ verifying the dihedral angles and geometry of the peptide bonds between the connecting residues (Figure S2). The energy minimized PDE5–EVO complex was used for subsequent molecular dynamics (MD) simulations and free energy calculations.

All-atom MD simulations of PDE5 in complex with the allosteric inhibitor EVO were started from the energy-minimized structure. The protein–inhibitor coordinates were extracted from this minimized structure and used to construct the MD simulation system under the same solvation and ionic conditions as described above. Prior to equilibration, the system was energy minimized for an additional 1000 steps using the conjugate gradient algorithm without any restraints to remove any steric clashes introduced during system setup. Next, we equilibrated the system for 20 ns by fixing the non-hydrogen atoms of the inhibitor. Three independent MD simulations were then carried out (each 1 *µ*s long) with configurations saved every 200 ps, during which no atoms were fixed.

We carried out all MD simulations using the CHARMM36 force field^40^ for the protein atoms and the CHARMM General Force Field (CGenFF) for the inhibitor atoms. ^41^ All simulations were conducted with a 2 fs integration time step in the isothermal–isobaric (NPT) ensemble. The temperature and pressure were maintained at 310 K and 1 bar, respectively, using the Langevin thermostat and the Nosé–Hoover barostat. We used periodic boundary conditions in all simulations and computed long-range electrostatic interactions using the Particle Mesh Ewald method.^42^ The van der Waals interactions were truncated at 12 Å, with a smooth switching function applied beyond 10 Å. We conducted all MD simulations using NAMD v3.0^38^ and the trajectories were analyzed using VMD.^36^

### Conformational analyses

Based on the data from MD simulations, we computed key metrics for the PDE5–EVO complex, including backbone root mean squared deviation (RMSD), per-residue C*_α_* root mean squared fluctuation (RMSF), and per-residue interaction energies between the PDE5 and EVO. We calculated the RMSD of the protein backbone atoms taking the energy minimized structure as a reference and the RMSF per residue was evaluated for the C*_α_* atom of each residue from its mean position. The interactions between the protein and the allosteric inhibitor were evaluated using the Protein-Ligand Interaction Profiler (PLIP) web tool. ^43^ We used the NAMDEnergy plugin in VMD^36^ to calculate the interaction energies between the PDE5 and the allosteric inhibitor complex. In these energy calculations, we used the force-field parameters from MD simulations, the nonbonded cutoff used in conducting MD simulations, as well as the Particle Mesh Ewald method for computing electrostatic interactions. The total interaction energy for each residue was determined as the sum of the electrostatic and van der Waals components. The PyMol^44^ and ChemAxon Marvin package^45^ were used for structural analysis and visualization.

### Alchemical Free Energy Calculations

We used alchemical free energy perturbation (FEP) calculations^46^ to quantify the relative binding free energy changes (ΔΔG) associated with residue perturbations in the PDE5-EVO complex. A total of 18 residue substitutions were evaluated (Table S2), comprising 11 alanine mutations, 4 single-residue mutations, 2 double-mutations, and a single triple-mutation, to assess their impacts on allosteric inhibitor binding. The modeled and energy minimized structure of the PDE5-EVO complex was used as the starting structure to ensure that interactions with the allosteric inhibitor are preserved. A thermodynamic cycle was constructed in which the vertical arms represent the physical binding of the inhibitor to wild type (WT) or mutated PDE5, and the horizontal arms represent the unphysical alchemical transformation of a wild type residue into its mutated form (Figure S3).^47,48^ Changes in free energy were calculated for the transformation in the inhibitor-bound complex and in the apo PDE5, with the relative change in binding free energy given by:

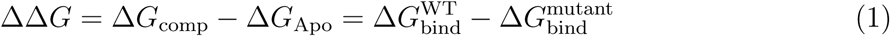

A dual-topology approach was used, with initial and final states present in the molecular topology and connected through the coupling parameter *λ* where *λ* = 0 corresponds to the WT, *λ* = 1 to the mutated, and intermediate values represent alchemical states. The free energy differences between the adjacent states were calculated using equation (2) and the total free energy change along the alchemical path was obtained by summing over the intermediate states, as described in equation (3):

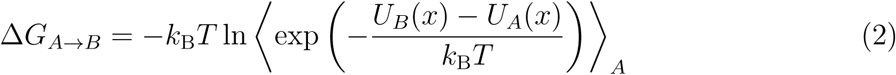

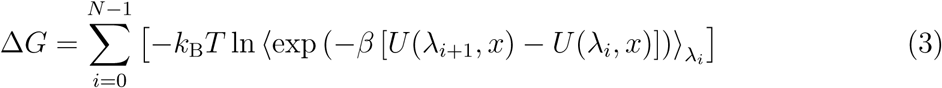

where *β* = 1*/k_B_T* (*k_B_* is the Boltzmann constant, *T* = 310 K) and *N* is the number of intermediate *λ* points. We set the value of “`alchDecouple`” to “`OFF`” in our free energy calculation protocol to scale the nonbonded interactions along the alchemical coordinate. This protocol allowed us to scale the nonbonded interactions of the mutated residue with their environment and within the mutated residue, which contributes to cumulative free energy. Furthermore, in our free energy calculations we set the “`alchElecLambdaStart`” to 0.5 and “`alchVdwLambdaEnd`” to 1.0 with “`alchVdWShiftCoeff`” value of 4.0. We used 25 equally spaced *λ* windows distributed over 50 ns of simulation time. We simulated each *λ* window for 2 ns and used the last 1.6 ns to estimate the free energy. We averaged the total free energy change (Δ*G*_comp_ and Δ*G*_free_) over forward and backward simulations and repeated them in triplicate with different initial velocities, resulting in ∼ 300 ns of simulation data per transformation (including both forward and backward) (Tables S3-S5). We performed all alchemical simulations using the same simulation parameters (force field, ensemble conditions, thermostat/barostat, and integration time step) as described for the MD simulations above.

We estimated the free energy differences using the bidirectional approach by incorporating samples from both forward and backward transformations. The related statistical error was calculated using the Bennet Acceptance Ratio (BAR) estimator implemented in the ParseFEP toolkit in VMD.^49^ To ensure convergence, we compared the underlying probability distributions that characterize the forward and backward transformations (Figures S4-S5). We reported the uncertainty in the averaged Δ*G*_comp_ and Δ*G*_Apo_ as the standard error of the mean (from three replicas) and the uncertainties in ΔΔ*G* were obtained by error propagation (combining individual uncertainties in quadrature, i.e., 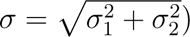.

## Results

### Allosteric Binding Pocket Comparison between PDE5 and PDE6 Isoforms

The crystal structure of the PDE5-EVO complex (PDB ID: 6VBI)^32^ contains an unresolved segment of the *α*14-helix and the M-loop within the allosteric pocket. As described in the methods section, we modeled and obtained a complete structure of the PDE5–EVO complex, with a proper backbone geometry and no Ramachandran outliers in the reconstructed region (Figure S2). The allosteric inhibitor also retained its crystallographic binding pose, preserving key interactions with the protein. Furthermore, to understand the residue-level differences in PDE5 and PDE6, we next performed the sequence and structural superposition of the catalytic domains of PDE5 and PDE6 (Figure S6), with particular emphasis on the allosteric pocket region (Figure 2). This comparison revealed that the majority of the residues that form the allosteric site are conserved between PDE5 and PDE6 or differ by conservative substitutions. The structural superposition further demonstrated a high degree of spatial overlap among secondary structure elements surrounding the allosteric pocket, indicating a largely preserved pocket architecture across these two enzymes.

**Figure 2:**
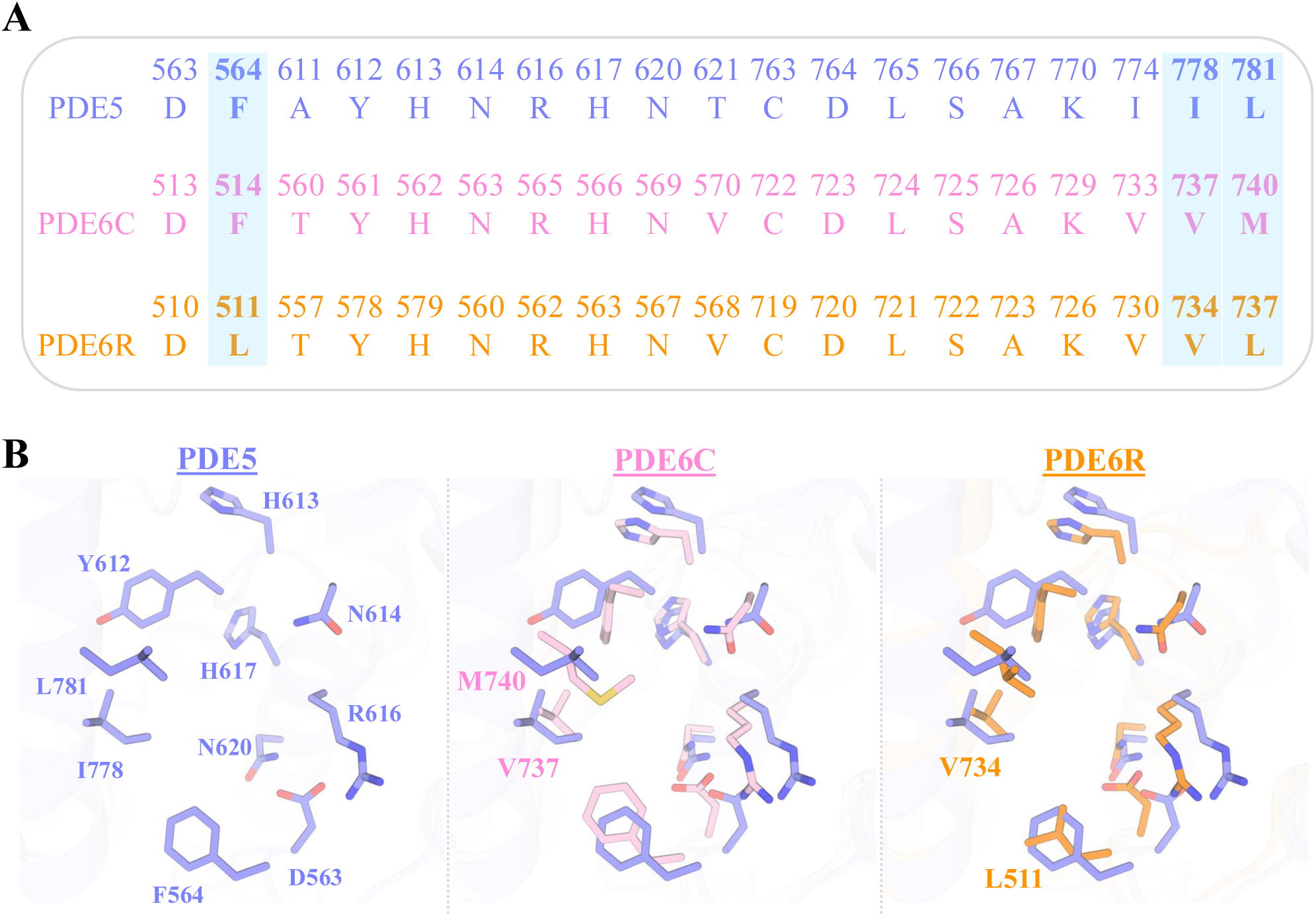
(A) Sequence alignment of PDE5 (UniProt ID: O76074) allosteric pocket residues with the corresponding positions in cone phosphodiesterase (PDE6C; UniProt ID: P51160) and rod phosphodiesterase (PDE6R; UniProt ID: P16499). Highlighted positions correspond to residues involved in EVO binding that differ in one or both PDE6 isoforms. (B) Structural superposition of the allosteric pocket regions of PDE5, PDE6C, and PDE6R. Each residue involved in inhibitor binding, identified from the PDE5–inhibitor complex (PDB ID: 6VBI), is shown in a stick representation.

Analysis of the crystal structure of the PDE5/EVO complex (PDB ID: 6VBI) showed that EVO interacts with residues D563, F564, Y612, H613, N614, R616, H617, N620, I778, and L781. A comparison with the cone and rod isoforms of PDE6 revealed differences at only two positions among the allosteric pocket residues of PDE5 involved in inhibitor binding (Figure 2B). In both PDE6 isoforms, I778 (PDE5) is replaced by V737 (PDE6C) and V734 (PDE6R) (Figure 2B). Additionally, L781 in PDE5 corresponds to M740 only in PDE6C, whereas PDE6R retains a leucine residue (L737). In contrast, PDE6R differs at position 564, where F564 in PDE5 is replaced by L511 (Figure 2B). Hence, residues that differed between PDE5 and PDE6 within the allosteric pocket were identified and selected for mutations in PDE5, enabling a systematic investigation of their contributions to the binding of the allosteric inhibitor.

### Structural and Energetic Evaluation of Binding Pocket Residues in PDE5

To elucidate the molecular determinants of inhibitor binding, we evaluated the structural and energetic contributions of key residues within the allosteric pocket of PDE5. We first assessed the stability of the PDE5-EVO complex by estimating the RMSD of the protein backbone atoms relative to the initial conformation. The calculated average RMSD for the PDE5 backbone atoms was 2.62 Å, reflecting that the protein structure was stable during the simulations, as it does not deviate significantly from the initial structure (Figure S7). The flexibility at the residue-level was assessed using the RMSF of the C*_α_* atoms of the residues. We observed RMSF values greater than 3 Å for the H-loop (residues 660-683) and M-loop regions (residues 798-812) (Figure S7). These regions in PDE5 are catalytically important structural elements.^27^ The other region with higher RMSF was a loop (742-748) connecting the *α*12 and *α*13 helices. In contrast, the RMSF values for the binding pocket residues were < 1.8 Å, reflecting the lower fluctuations observed for the residues in the binding site (Figure S7). Given that loop regions are known to have a higher flexibility, the RMSF for residues in various loops was higher compared to residues on helices or other secondary structure motifs.

In addition, inspection of the binding pocket and the binding mode of the inhibitor revealed that different chemical moieties of EVO engage specific residues through complementary interactions within the allosteric pocket, thereby contributing to the overall binding affinity. The indole ring of the inhibitor forms stabilizing hydrogen bonds with D563 (D563-O*δ*1· · · N2-EVO; 3.06 Å) and N614 (N614-N*δ*2· · · O3-EVO; 3.14 Å), anchoring the inhibitor within the pocket, while additional interactions in the pocket involve residues F564 and R616, which further support the positioning of the inhibitor through hydrophobic contacts (Figure 3B). The methoxy substituent at the indole ring establishes a hydrogen bond with N614, reinforcing stabilization of the ligand. Within the hydrophobic core of the pocket, the quinazolinone core is stabilized primarily through van der Waals contacts with A767, N620 and K770. Collectively, these interactions demonstrate that EVO utilizes a combination of polar and hydrophobic contacts, enabling optimal accommodation and stabilization within the allosteric site of PDE5.

**Figure 3:**
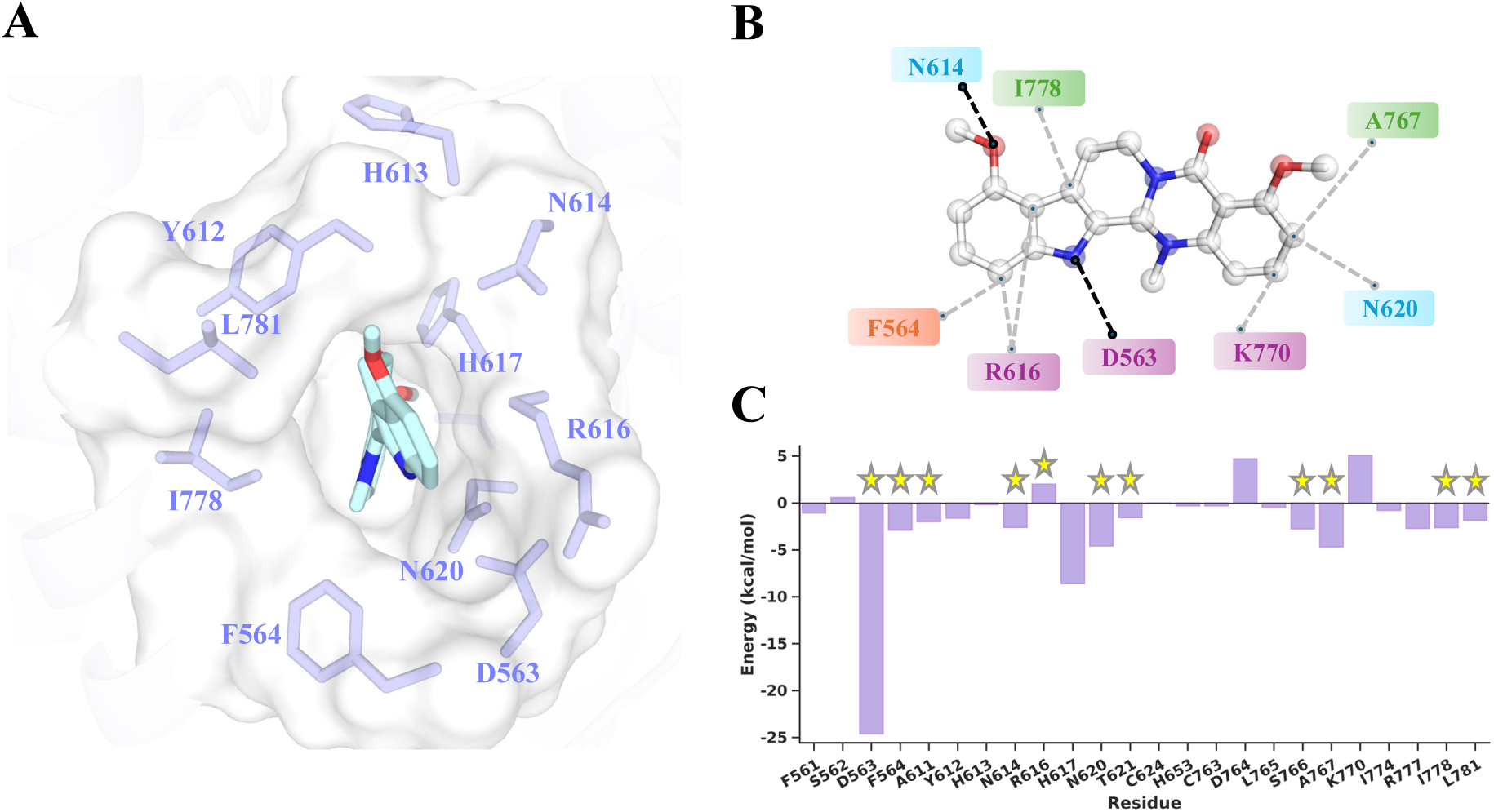
(A) Allosteric binding pocket of PDE5 in complex with EVO (PDB ID: 6VBI). (B) A two-dimensional interaction map of EVO with residues in the PDE5 allosteric pocket. The hydrogen bonding interactions are depicted as black dashed lines, while hydrophobic interactions are shown as gray dashed lines. The residues are classified according to their physicochemical properties as polar (blue), non-polar aliphatic (green), non-polar aromatic (orange), and charged (purple). (C) Non-bonded interaction energy between all atoms of each binding pocket residue and the inhibitor. An asterisk marks those residues that are selected for further mutations.

We next calculated per-residue non-bonded interaction energies from MD trajectories to initially assess the energetic contributions of individual residues in binding of EVO. The residue D563 showed the highest energy contribution with a non-bonded interaction energy of -24.60 kcal/mol (Figure 3C). Additional residues with significant energetic contributions included F564, N614, H617, N620, S766, A767 and I778 (Figure 3C). To systematically select residues for mutations, we identified residues that contribute substantially to EVO binding, specifically those with per-residue interaction energies lower than -2.00 kcal/mol. We also considered pocket residues that differ in sequence relative to the corresponding positions in PDE6 isoforms. Based on these criteria, we identified D563, F564, A611, N614, R616, N620, T621, S766, A767, I778, and L781 as candidates for alanine substitution. The H617 residue was not included due to its variable protonation and tautomeric states. Overall, these calculations allowed us to initially assess the interaction energies of individual residues for further assessment via subsequent alchemical FEP calculations.

### Alanine-scanning of Selected Binding Pocket Residues

We first performed alchemical free energy calculations by transforming each selected residue into an alanine residue. If the wild-type residue was an alanine, it was mutated to a serine residue (smallest polar residue). A total of 11 residues (D563, F564, A611, N614, R616, N620, T621, S766, A767, I778, and L781) within the binding pocket were individually mutated, and the resulting changes in the binding free energy (ΔΔG) were calculated (Figure 4A). The ΔΔG values reflected a spectrum of effects, ranging from strong stabilization to significant destabilization, highlighting their differing energetic contributions to inhibitor binding (Table S3). To interpret these results, we applied a classification scheme based on the magnitude of ΔΔG: mutations with ΔΔG > +1.0 kcal/mol were considered strongly destabilizing, ΔΔG = +0.3 to +1.0 kcal/mol moderately destabilizing, ΔΔG = –0.3 to +0.3 kcal/mol neutral, ΔΔG = –0.3 to –1.0 kcal/mol moderately stabilizing, and ΔΔG < –1.0 kcal/mol strongly stabilizing.

**Figure 4:**
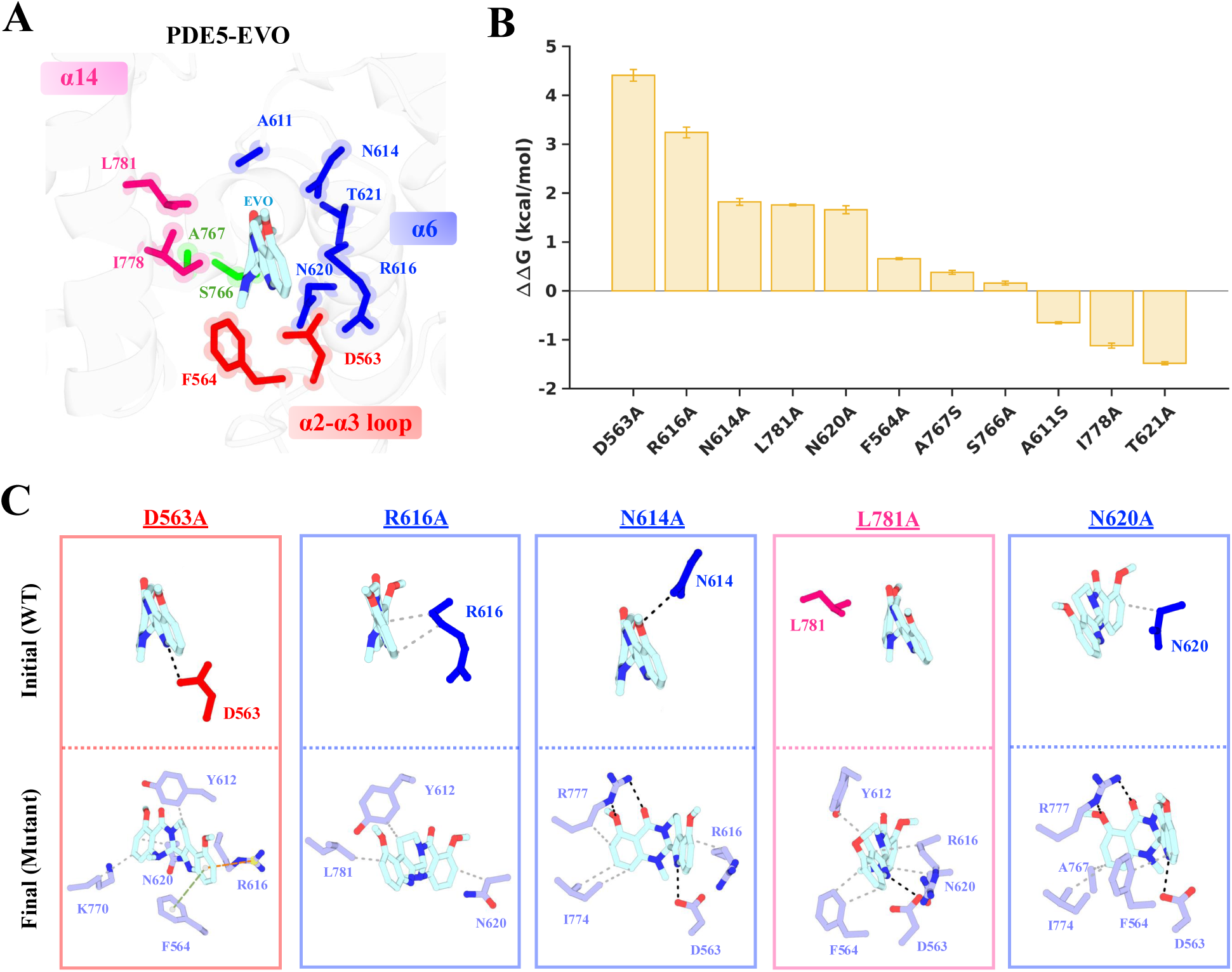
Energetics of alanine substitutions in PDE5 and associated structural changes in the allosteric inhibitor EVO-PDE5 complex. (A) The structure of the PDE5–EVO complex highlighting residues selected for alanine substitution. The residues are colored according to their positions within the secondary structure elements: *α*14-helix (pink), *α*6-helix (blue), *α*2-*α*3 loop (red), and the *α*14-*α*13 loop (green). (B) Shown are relative binding free energy difference (ΔΔG) for mutations of each residue to alanine. (C) Snapshots highlighting the interactions between EVO and selected residues (D563, R616, N614, L781, N620) in the PDE5 complex. Upper panels show only the initial WT residue interactions with EVO prior to alanine substitution, and lower panels show the rearranged interactions following the mutations.

Using these criteria, the strongly destabilizing mutations included D563A (+4.41 kcal/mol), R616A (+3.24 kcal/mol), L781A (+1.76 kcal/mol), N620A (+1.66 kcal/mol), and N614A (+1.82 kcal/mol), indicating that these residues are critical to binding of EVO (Figure 4A). The mutations F564A (+0.66 kcal/mol) and A767S (+0.38 kcal/mol) resulted in moderate destabilization, while S766A (+0.16 kcal/mol) was essentially neutral, having minimal effect on inhibitor affinity (Figure 4A). In contrast, T621A (–1.48 kcal/mol), I778A (–1.12 kcal/mol), and A611S (–0.65 kcal/mol) were stabilizing, suggesting that subtle modifications in the size or polarity of a given sidechain may result in local flexibility or minor rearrangements in the allosteric pocket, favoring EVO binding.

We further examined how each substitution of alanine altered protein–inhibitor interactions, thus providing a mechanistic rationale for the observed changes in binding free energy (Figure 4C, lower panel). The residue D563 serves as a key hydrogen-bond anchor for EVO, and its substitution to alanine abolished all hydrogen bonding interactions in the pocket (Figure 4C). The R616A mutation disrupted interactions with F564 along with disrupting the hydrogen-bond network involving residues D563 and N614. This mutation resulted in EVO forming weaker hydrophobic interactions with the residues N620 and Y612, ultimately destabilizing the protein-inhibitor complex. The mutation N614A abolished the hydrogen bond between the amide nitrogen N*δ*2 of N614 and the O3 atom of EVO (Figure 4C and Figure S8). As a result, EVO shifted towards the *α*14-helix and formed compensatory interactions with the residues R777 and I774. The L781A mutation mainly affected the hydrophobic packing at the distal end of the binding pocket (interactions with quinazolinone core).

Although the inhibitor largely retained its orientation, hydrophobic contacts with residues K770, I778, and A767 were lost. Upon N620A mutation, the hydrophobic contact between N620 and EVO was lost, and the inhibitor was reoriented toward the *α*14-helix, interacting with R777 and I774. Interestingly, I778A slightly stabilized the complex. By replacing a bulky hydrophobic residue with alanine, this mutation decreased steric hindrance, allowing EVO to adopt a more favorable binding conformation and strengthening the D563 hydrogen bond (D563-O*δ*1···N2-EVO; 2.8 Å). Similarly, T621A and A611S promoted subtle rearrangements that favored hydrogen-bonding and hydrophobic interactions, highlighting how local pocket flexibility can enhance inhibitor accommodation. Collectively, these results identified D563, N614, R616, N620, and L781 as critical residues for EVO binding, while residues I778, T621, and A611 contributed to pocket adaptability, allowing the inhibitor to explore alternative stable conformations without a substantial loss in affinity. Thus, the alanine scanning free energy calculations revealed a hierarchy in interactions of pocket residues with the allosteric inhibitor.

### Effect of PDE6 Derived Single Mutations

Following a systematic evaluation of alanine mutations of residues critical to EVO binding in PDE5, we investigated the effect of homolog specific residue-level differences between PDE5 and PDE6 (cone and rod) on inhibitor recognition. We performed alchemical transformations on four PDE5 residues: F564, R616, I778, and L781, which interact directly with EVO and differ from their corresponding residues in PDE6 isoforms (PDE6C and PDE6R). These substitutions, hereafter referred to as PDE6 derived mutations, were used to mimic the sequence variations found in PDE6 and to probe their impacts on the allosteric pocket and EVO binding. This approach enables a comparative evaluation of EVO binding between PDE5 and PDE6 and facilitates the identification of residues that govern selectivity.

The calculated changes in binding free energy (ΔΔG) for single residue substitutions are shown in Figure 5A and summarized in Table S4. The ΔΔG values range between –0.69 and +1.49 kcal/mol, indicating effects spanning marginal stabilization to significant destabilization (Figure 5A). The R616Q transformation exhibited the strongest destabilizing effect (ΔΔG = +1.49 kcal/mol) by abolishing all hydrogen bonds with EVO. In this altered binding mode, the inhibitor primarily engaged with L781 and I778 through hydrophobic contacts, suggesting that simultaneous mutations in R616 and L781/I778 would likely have a synergistic destabilizing effect. The mutation F564L induced a moderate destabilization (ΔΔG = +0.50 kcal/mol), primarily through loss of hydrophobic interaction with EVO at the position 564, while the remainder of the residue contacts remained largely unperturbed (Figure 5B). On the other hand, L781M led to a marginal destabilization (ΔΔG = +0.41 kcal/mol) effect. Although the inhibitor was retained within the pocket, the critical hydrogen bonding interactions of EVO with D563 and N614 were lost and compensatory interactions formed with neighboring residues in the *α*14-helix. In contrast, the I778V substitution resulted in slight stabilization (ΔΔG = –0.69 kcal/mol), enabling the inhibitor to adopt a conformation that forms additional hydrogen bonds with R777 (R777-N*ε*· · · O1-EVO; 2.7 Å and R777-N*η*2· · · O1-EVO; 2.9 Å) and L566 (L566-O· · · N2-EVO; 2.9 Å), highlighting the role of local sidechain geometry in modulating pocket flexibility. Collectively, these PDE6-derived mutations demonstrated that R616 and F564 are key determinants of EVO binding, whereas L781 and I778 predominantly modulate pocket flexibility and inhibitor orientation. These observations provide mechanistic insights into how homolog specific residue differences between PDE5 and PDE6 influence inhibitor recognition and allosteric pocket dynamics, complementing the residue-level mapping obtained from the alanine mutations in PDE5.

**Figure 5:**
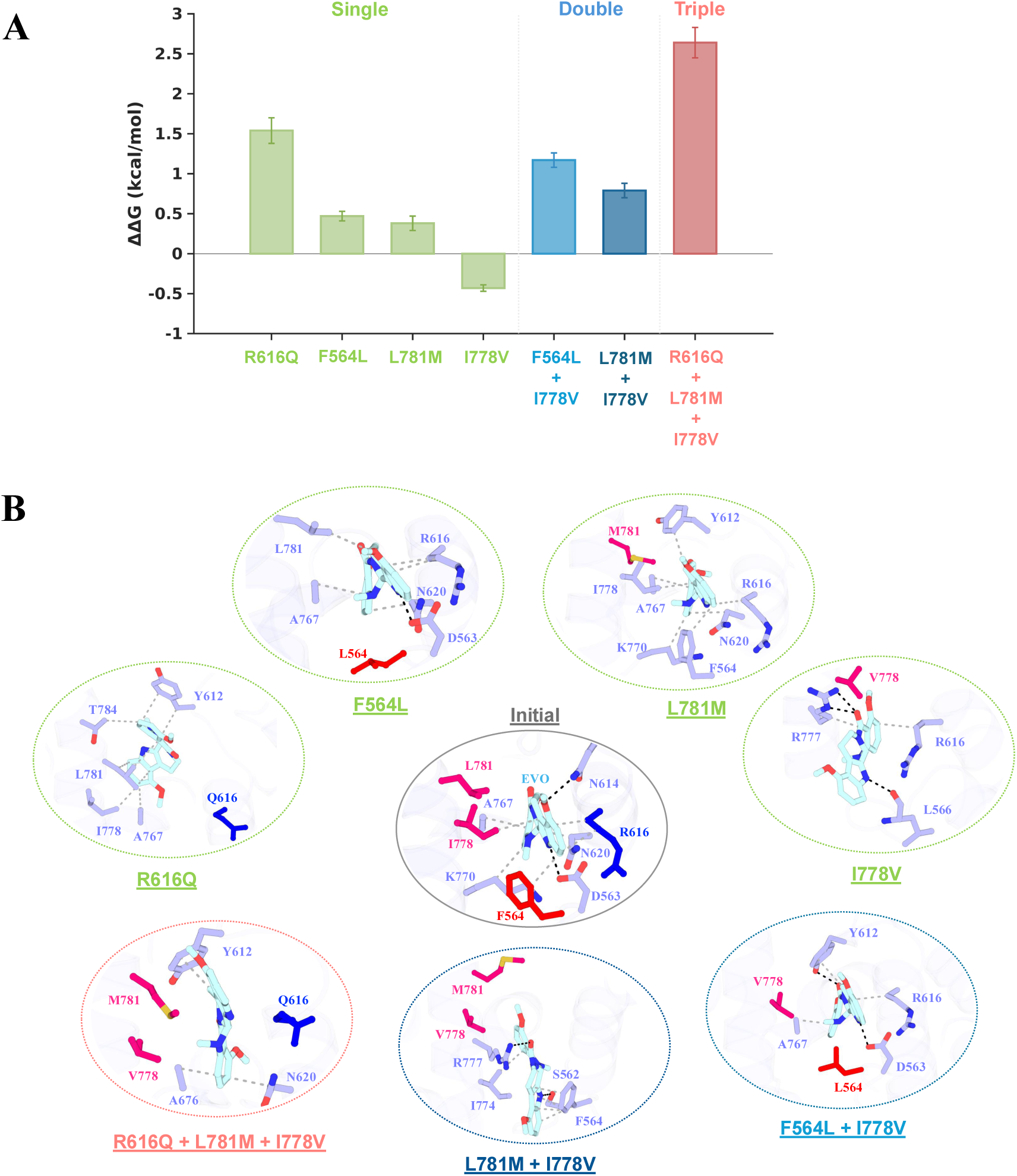
Energetic and structural effects of PDE6-derived residue substitutions on evodiamine (EVO) binding in PDE5. (A) Relative binding free energy changes (ΔΔG) obtained from alchemical free-energy calculations for PDE6 derived substitutions introduced into the PDE5 allosteric pocket. The mutations are grouped as single (R616Q, F564L, L781M, I778V), double (I778V+F564L, mimicking PDE6R and I778V+L781M, mimicking PDE6C), and triple (I778V+L781M+R616Q, corresponding to a PDE6 variant previously studied^32^). (B) Snapshots showing interactions between residues and the inhibitor within the PDE5-EVO complex for various PDE6-derived mutations. Hydrogen bonding interactions are depicted as black dashed lines, while hydrophobic interactions are shown as grey dashed lines. The mutated residues are colored according to their position within defined secondary structural elements: *α*14 helix (pink), *α*6-helix (blue), *α*2-*α*3 loop (red) (see figure 4).

### PDE6 Isoform Specific Double and Triple Mutations Reveal Selectivity Determinants

We further evaluated homolog specific residue pairs to assess their combined effects by modeling the allosteric pocket environments of the PDE6 cone and rod isoforms. Specifically, we constructed the I778V + L781M combination to mimic the PDE6 cone variant and I778V + F564L to mimic the PDE6 rod variant. In addition, an R616Q substitution was introduced to generate the triple mutant I778V + L781M + R616Q, corresponding to the PDE6 cone variant examined in a previous experimental study.^32^ These combinations allowed us to evaluate how combinatorial residue changes influence EVO binding and the conformational dynamics of the allosteric pocket. The calculated ΔΔG values for these double/triple mutations are shown in Figure 5A and summarized in Table S5. The PDE6C double mutant exhibited a moderate destabilization with a ΔΔG value of +0.71 kcal/mol (Figure 5A). In contrast, the PDE6R double mutant resulted in a stronger destabilizing effect with a ΔΔG value +1.12 kcal/mol, indicating that residue interactions in the rod variant more substantially perturb the binding of allosteric inhibitor (Figure 5A). Furthermore, the triple mutant corresponding to the PDE6C experimental variant resulted in an even stronger destabilization, with a ΔΔG value of +2.29 kcal/mol, reflecting a stronger impact on inhibitor stability within the allosteric pocket (Figure 5A).

Structural analysis further confirmed these observations, as distinct mechanistic consequences were observed for these combined substitutions. In PDE6R (F564L and I778V) the mutations involved residues both in the *α*14-helix (I778V) and the *α*2-*α*3 loop (F564L). Although the hydrogen bond between the residue D563 and EVO is retained, there is a significant loss of hydrophobic interactions with EVO, accounting for the stronger destabilization observed in this variant (Figure 5B). In contrast, in PDE6C (L781M and I778V), both residues are located within the *α*14-helix, and their combined effect alters the EVO conformation. The inhibitor establishes a new hydrogen bond between the carbonyl oxygen of the quinazolinone moiety and R777 (R777-N*η*1· · · O1-EVO; 3.5 Å), while the indole ring is stabilized through multiple interactions with F564 and a hydrogen bond with S562 (S562-O*γ*· · · N2-EVO; 3.5 Å) (Figure 5B). Furthermore, the PDE6C triple mutation (I778V, L781M, and R616Q) carrying the experimentally reported variant (R616Q) disrupted the hydrogen bonding network of EVO. Specifically, the R616Q substitution disrupted key interactions with F564, D563, and N614, leading to a conformational shift of the inhibitor that abolishes hydrogen-bonding interactions with D563 and N614 (Figure 5B). These results indicate that coordinated residue changes can have additive or synergistic effects on inhibitor binding, with the PDE6R substitutions eliciting a stronger destabilizing effect than the PDE6C variant. Our results highlight the critical role of specific residues in modulating EVO conformation and provide mechanistic insights into how sequence variations between PDE5 and PDE6 may differentially influence allosteric inhibitor recognition.

## Discussion

Selective inhibition of PDEs remains challenging due to high sequence conservation among their catalytic domains.^50^ Current orthosteric inhibitors of PDE5 such as sildenafil and tadalafil target the conserved catalytic pocket and often resulting in off-target activity against PDE6, thus motivating alternative strategies that exploit less conserved regulatory regions.^32,33,51,52^ The identification of a unique allosteric pocket in PDE5 and the discovery of evodiamine derived inhibitors that selectively bind to this site represent key advances toward achieving isoform selectivity. ^32^ In this study, we employed an integrated computational framework combining structural modeling, all-atom simulations, and alchemical free energy calculations to elucidate the molecular determinants of allosteric inhibitor recognition in PDE5 and to rationalize how sequence variation in PDE6 isoforms modulate inhibitor binding. By systematically analyzing single, double, and triple residue mutation, our results provide a detailed residue-level characterization of the PDE5 allosteric pocket and elucidate the mechanistic basis for differential inhibitor recognition between PDE5 and PDE6.

Together, the per-residue non-bonded interaction energy decomposition and alanine scanning free energy calculations identified a small subset of residues (D563, N614, R616, N620, and L781) as dominant contributors to the binding of EVO. These residues form an interconnected network of hydrogen bonds and hydrophobic contacts that anchor EVO and restrict its conformational flexibility. Among these, D563 is a key anchoring residue, contributing the highest favorable interaction energy and consistently maintaining a stabilizing hydrogen bond with EVO in all simulations. The alanine substitution at D563 resulted in strongest destabilizing effects (+4.41 kcal/mol), highlighting the critical role of hydrogen bonding and electrostatic interactions in stabilizing EVO within the pocket. A previous study^32^ that characterized EVO derivatives as PDE5 allosteric inhibitors identified conserved acidic residues within the allosteric pocket as key anchoring points for ligand binding. Consistent with these findings, our results indicate that the acidic residue D563 serves as the central interaction hotspot governing inhibitor stabilization.

Beyond D563, two other residues, N614 and R616, form critical interactions with EVO within the allosteric pocket. The disruption of these interactions, either through alanine substitution or PDE6-derived substitutions, led to pronounced destabilization and conformational rearrangements of EVO. In contrast, mutations such as I778A, T621A, and A611S resulted in modest stabilization, suggesting that a partial reduction in sidechain steric hindrance or polarity can enhance local flexibility and allow EVO to adopt more favorable binding conformations. These findings emphasize that the binding affinity may not be solely dictated by strong anchoring interactions, but also by a balance between rigidity and adaptability within the pocket. Notably, increased local flexibility or reduced steric hindrance can enhance binding by allowing the ligand to adopt a more favorable conformation in the allosteric pocket.

Moreover, probing the impact of homolog-specific substitutions further highlighted residues contributing to EVO selectivity for PDE5 over PDE6. Single PDE6-derived mutations revealed that R616Q and F564L are strongly destabilizing, whereas I778V slightly stabilizes EVO binding. Our findings suggest that residues F564 and R616 are major determinants of binding affinity, whereas substitutions at I778 and L781 primarily influence pocket geometry and ligand orientation. Notably, the R616Q substitution derived from a previously reported PDE6 variant^32^ abolished all hydrogen bonding with EVO and the inhibitor predominantly engaged in hydrophobic interactions in the allosteric pocket. Although the residue R616 is an important modulator of allosteric inhibitor binding, it should be noted that the R616Q substitution does not represent a naturally occurring residue variation between PDE5 and native PDE6 isoforms, but was introduced to assess a PDE6-variant reported in an experimental study.^32^ In this specific context, the loss of R616-mediated interactions led to a substantial destabilization of EVO binding, indicating that the inhibitor engages with this residue to stabilize a PDE5-like allosteric environment. Consequently, inhibitors designed to form favorable interactions with R616 may enhance PDE5 selectivity, whereas inhibitor chemotypes that are less dependent on this interaction may exhibit broader cross-reactivity. In contrast, the I778V substitution slightly stabilized EVO binding by enabling alternative hydrogen bonding with R777, illustrating how subtle differences in the size of a sidechain (smaller Val instead of Ile) can reshape the local interaction pattern without compromising the affinity.

Furthermore, the combined substitutions designed to replicate the allosteric site of PDE6 isoforms revealed clear differences in their effects on EVO binding. Mutations derived from the PDE6C isoform (I778V + L781M) are conservative and produce moderate destabilization, indicating that modifications confined to the *α*14-helix primarily influence inhibitor orientation while largely preserving the hydrogen-bond network. However, double mutations reflecting PDE6R (I778V + F564L) span both the *α*14-helix and the *α*2-*α*3 loop, resulting in a more pronounced change in the hydrophobic packing due to the loss of key interactions with two residues in the pocket (F564 and I778). Our findings indicate that the differential impact on EVO binding arises from the spatial distribution of these substitutions within the allosteric pocket. Furthermore, the triple mutant (I778V + L781M + R616Q) exhibited the highest effect on the binding affinity, reflecting a synergistic perturbation of helix packing and key hydrogen bonding interactions. This substantial destabilization is consistent with the effects observed for individual substitution of R616Q, where loss of key hydrogen bonds with EVO redirected interactions toward the *α*14-helix, particularly involving residues T784, L781 and I778. In the triple mutant, this effect is amplified by additional mutations (I778V and L781M), which collectively disrupt interactions with residues in the *α*14-helix and hydrogen bonding interactions involving D563 and N614, leading to substantial reorganization of the inhibitor within the allosteric pocket. These findings highlight how specific substitutions modulate interactions in the allosteric pocket and provide a mechanistic basis for selectivity differences between the allosteric sites of PDE5 and PDE6.

Overall, our free energy calculations indicate that the binding of EVO to PDE5 is governed by a modular interaction pattern in which distinct chemical moieties of the inhibitor engage with complementary regions of the allosteric pocket through coordinated polar and hydrophobic interactions. The indole ring, a key heterocycle in the EVO core (Figure 1B) anchors the inhibitor by forming hydrogen bonds with D563 and N614, consistent with the strong destabilization observed upon alanine substitutions of these residues. These results are in agreement with previous structural studies identifying the indole moiety as a key pharmacophoric element^53^ that orients EVO within the allosteric cavity and modulates the conformation of the H-loop.^27^ The fused pyridine ring serves as an anchor for the indole moiety and the quinazolinone core, indicating a role in optimizing ligand positioning rather than serving as a primary affinity determinant. Deeper within the allosteric pocket, the quinazolinone core engages residues from the *α*6-helix as well as the *α*14-helix, highlighting stabilization through hydrogen-bonds and van der Waals interactions. Additionally, it can form compensatory interactions, including alternative hydrogen bonding with R777 and Y612 in PDE6 derived variants, highlighting the adaptability of this region to local pocket variations. The quinazolinone fused methoxybenzene contributes additional hydrophobic contacts that stabilize EVO in the hydrophobic core of the allosteric pocket. Although these interactions are individually less energetically dominant than the polar interactions of the indole moiety, they help to rigidify the overall scaffold orientation and provide a cumulative stabilizing effect. Consistent with this architecture, a small aliphatic residue at position 778 favors the accommodation of the inhibitor, as reflected by a modest stabilization upon substitutions of I778V and I778A. Alanine substitutions at F564 and L781 are more destabilizing than conservative mutations (F564L and L781M), highlighting the importance of steric and hydrophobic balance at these positions. Collectively, residues from the *α*6-helix and the *α*2-*α*3 loop emerge as key contributors to the binding of the allosteric inhibitor to the PDE5 allosteric pocket. Whereas, conservative variations within the *α*14-helix had minor effects, indicating that this region tolerates small structural perturbations without a substantial loss in the binding affinity of the inhibitor.

Furthermore, our findings establish structure–affinity correlations that can inform the rational design of selective allosteric modulators for PDE5. The crucial role of the indole–D563/N614 hydrogen bonding network suggests that incorporating or enhancing polar groups capable of maintaining these interactions should be prioritized in lead optimization. The modulatory role of quinazolinone in engaging core hydrophobic and polar contacts highlights an opportunity to optimize ring substituents that either reinforce these interactions or introduce additional favorable contacts, particularly with residues such as R777 that may become more accessible in certain conformational states. Meanwhile, modifications to fused methoxybenzene that strengthen the *π*-stacking interactions and enhance the hydrogen bonding within the pocket core, such as the introduction of a hydroxyl group, may improve binding potency and selectivity. Hence, the findings of our study collectively highlight several key principles relevant to drug design. First, multi-moiety inhibitors such as evodiamine can achieve high affinity and selectivity by simultaneously engaging multiple complementary residues within the allosteric pocket. Second, the balance between anchor residues (e.g., D563, R616) and flexible or adaptable residues (e.g., I778, *α*14-helix) governs pocket tolerance and inhibitor accommodation. Third, isoform-specific variations, as seen in PDE6C versus PDE6R, can modulate inhibitor binding through subtle shifts in hydrogen bonding and hydrophobic interactions, providing a structural basis for selective inhibition. These insights complement existing experimental data and inform the structure based design of selective PDE inhibitors.

## Conclusion

In summary, our study revealed the molecular basis for allosteric inhibition of PDE5 by integrating structural and energetic evaluation of the inhibitor-bound allosteric pocket. By comprehensive free energy calculations, we identified key determinants of EVO binding and revealed how sequence variations in PDE6 isoforms differentially perturb allosteric pocket interactions. In the PDE5 allosteric pocket, EVO participates in a network of interactions distributed across multiple structural regions, involving residues from distinct *α*-helices and loop segments. Strong anchoring interactions involve D563 and N614 (hydrogen bonding) and R616 (polar and hydrophobic contacts), while adaptable peripheral residues such as F564, A611, I778, L781, and positions within the *α*14-helix (K770, R777, I774) collectively contribute to pocket flexibility and accommodate inhibitor conformations. Our results highlight that distinct chemical moieties of EVO interact with complementary regions of the pocket, collectively stabilizing the inhibitor through polar and hydrophobic contacts. Furthermore, the evaluation of PDE6-derived mutations revealed isoform specific effects, with rod-PDE6 variations inducing a stronger destabilizing effect than cone-PDE6 substitutions. Collectively, our study reveals mechanistic insights into allosteric inhibition of PDE5 and informs rational strategies for the design of selective inhibitors.

## Acknowledgement

We acknowledge the financial support provided by the National Institutes of Health (NIH) through Grant R35GM138217. The content is solely the responsibility of the authors and does not necessarily represent the official views of the NIH. We are grateful for computational support through Premise, a central shared HPC cluster at UNH supported by the Research Computing Center.

## Data Availability Statement

All methodological details for the data generated in this study are available in the Materials and Methods section including PDB codes of input files, simulation parameters, and error analysis. The input files for conventional MD simulations and FEP calculations for mutations in the PDE5-EVO complex are available at github: (https://github.com/jyoti-combio/PDE5_JCIM.git). The free energy data averaged over forward and backward transformations for each mutation in complex and solution are also provided (Tables S3–S5). The simulation and visualization softwares and the force field used to carry out calculations are openly available.

