## Supporting Information for "Molecular Determinants of Allosteric Inhibitor Affinity and Selectivity in PDE5"

**Table S1:** The residues in the allosteric pocket of PDE5 and the corresponding residue identities and positions in PDE6C (cone) and PDE6R (rod). Each amino acid is denoted using the single-letter code. Sequence information was obtained from UniProt entries O76074 (PDE5), P51160 (PDE6C), and P16499 (PDE6R).

| PDE5 Position | PDE5 | PDE6C Position | PDE6C | PDE6R Position | PDE6R |
|---|---|---|---|---|---|
| **563** | D | 513 | D | 510 | D |
| **564** | F | 514 | F | 511 | L |
| **611** | A | 560 | T | 557 | T |
| **612** | Y | 561 | Y | 578 | Y |
| **613** | H | 562 | H | 579 | H |
| **614** | N | 563 | N | 560 | N |
| **616** | R | 565 | R | 562 | R |
| **617** | H | 566 | H | 563 | H |
| **620** | N | 569 | N | 567 | N |
| **621** | T | 570 | V | 568 | V |
| **763** | C | 722 | C | 719 | C |
| **764** | D | 723 | D | 720 | D |
| **765** | L | 724 | L | 721 | L |
| **766** | S | 725 | S | 722 | S |
| **767** | A | 726 | A | 723 | A |
| **770** | K | 729 | K | 726 | K |
| **774** | I | 733 | V | 730 | V |
| **778** | I | 737 | V | 734 | V |
| **781** | L | 740 | M | 737 | L |

**Table S2:** Summary of mutations performed in PDE5 and their corresponding residues in PDE6 isoforms. All substitutions were introduced and simulated exclusively in the PDE5 structure. The corresponding residues in PDE6C (cone) and PDE6R (rod) are shown for sequence mapping and comparative interpretation only; no mutations were performed directly in PDE6 isoforms.

| PDE5 Mutation (Performed) | Mutation Type | PDE5 Residue (Mutated) | PDE6C (Cone) Corresponding Residue | PDE6R (Rod) Corresponding Residue | Substitution rationale |
|---|---|---|---|---|---|
| **D563A** | Single (Alanine scan) | D563 → A | D513 | D510 | Alanine substitution at a fully conserved position |
| **F564A** | Single (Alanine scan) | F564 → A | F514 | L511 | Alanine substitution at an aromatic position with rod-specific variation |
| **A611S** | Single | A611 → S | T560 | T557 | Substitution from non-polar to polar |
| **N614A** | Single (Alanine scan) | N614 → A | N563 | N560 | Alanine substitution of a conserved polar residue |
| **R616A** | Single (Alanine scan) | R616 → A | R565 | R562 | Alanine substitution of a conserved charged residue |
| **N620A** | Single (Alanine scan) | N620 → A | N569 | N567 | Alanine substitution of a conserved polar residue |
| **T621A** | Single (Alanine scan) | T621 → A | V570 | V568 | Alanine substitution at a position varying in PDE6 isoforms |
| **S766A** | Single (Alanine scan) | S766 → A | S725 | S722 | Alanine substitution of a conserved residue |
| **A767S** | Single | A767 → S | A726 | A723 | Substitution from non-polar to polar |
| **I778A** | Single (Alanine scan) | I778 → A | V737 | V734 | Alanine substitution at a position varying in PDE6 isoforms |
| **L781A** | Single (Alanine scan) | L781 → A | M740 | L737 | Alanine substitution at a position with cone-specific variation |
| **F564L** | Single (Non-alanine) | F564 → L | F514 | L511 | Rod-mimicking substitution |
| **R616Q*** | Single | R616 → Q | R565 | R562 | Cone-mimicking (adapted from experimental study) |
| **I778V** | Single (Conservative) | I778 → V | V737 | V734 | PDE6 Isoform-matched conservative substitution |
| **L781M** | Single (Conservative) | L781 → M | M740 | L737 | Cone-mimicking substitution |
| **I778V + L781M** | Double | I778 → V + L781 → M | V737 + M740 | V734 + L737 | Cone-like combined substitutions |
| **I778V + F564L** | Double | I778 → V + F564 → L | V737 + F514 | V734 + L511 | Rod-like combined substitutions |
| **I778V + L781M + R616Q*** | Triple | I778 → V + L781 → M + R616 → Q | V737 + M740 + R565 | V734 + L737 + R562 | Cone-like triple substitution |

*substitution derived from a PDE6 variant used in the experimental study reporting evodiamine derivatives as PDE5 allosteric inhibitors.

**Table S3:** Free energy perturbation data for amino-acid mutations in PDE5 when bound to the inhibitor (ΔG Comp) and unbound in solution (ΔG Apo). We estimated the free energy differences using the bidirectional approach by incorporating samples from both forward and reverse alchemical transformations. The relative binding free energy (ΔΔG) is reported for each mutation.

| Mutation | Run | Sim. length | ΔG Comp | ΔG Apo | ΔΔG (kcal/mol) |
|---|---|---|---|---|---|
| **D563A** | Run 1 | 50 ns | 130.23 ± 0.15 | 123.59 ± 0.14 | **4.41 ± 0.12** |
| | Run 2 | 50 ns | 128.43 ± 0.21 | 125.39 ± 0.15 | |
| | Run 3 | 50 ns | 128.71 ± 0.15 | 125.15 ± 0.13 | |
| | **Average** | | **129.12 ± 0.10** | **124.71 ± 0.08** | |
| **F564A** | Run 1 | 50 ns | -6.49 ± 0.05 | -7.71 ± 0.02 | **0.66 ± 0.02** |
| | Run 2 | 50 ns | -6.95 ± 0.05 | -7.23 ± 0.03 | |
| | Run 3 | 50 ns | -7.22 ± 0.03 | -7.70 ± 0.02 | |
| | **Average** | | **-6.88 ± 0.02** | **-7.54 ± 0.01** | |
| **A611S** | Run 1 | 50 ns | 3.28 ± 0.04 | 4.48± 0.04 | **-0.65 ± 0.02** |
| | Run 2 | 50 ns | 3.84 ± 0.05 | 4.33 ± 0.03 | |
| | Run 3 | 50 ns | 4.03 ± 0.04 | 4.29 ± 0.06 | |
| | **Average** | | **3.71 ± 0.02** | **4.36 ± 0.02** | |
| **N614A** | Run 1 | 50 ns | 78.57 ± 0.06 | 74.97 ± 0.11 | **1.82 ± 0.07** |
| | Run 2 | 50 ns | 77.81 ± 0.11 | 76.64 ± 0.10 | |
| | Run 3 | 50 ns | 78.19 ± 0.09 | 77.50 ± 0.07 | |
| | **Average** | | **78.19 ± 0.05** | **76.37 ± 0.05** | |
| **R616A** | Run 1 | 50 ns | 267.75 ± 0.16 | 264.30 ± 0.12 | **3.24 ± 0.11** |
| | Run 2 | 50 ns | 268.25 ± 0.15 | 265.30 ± 0.13 | |
| | Run 3 | 50 ns | 268.49 ± 0.12 | 265.17 ± 0.14 | |
| | **Average** | | **268.16 ± 0.08** | **264.92 ± 0.08** | |
| **N620A** | Run 1 | 50 ns | 79.11 ± 0.09 | 77.26 ± 0.10 | **1.66 ± 0.08** |
| | Run 2 | 50 ns | 78.56 ± 0.12 | 77.61 ± 0.08 | |

| | | | | | |
|---|---|---|---|---|---|
| | Run 3 | 50 ns | 78.33 ± 0.11 | 76.15 ± 0.08 | |
| | **Average** | | **78.66 ± 0.06** | **77.00 ± 0.05** | |
| **T621A** | Run 1 | 50 ns | 17.45 ± 0.05 | 19.12 ± 0.05 | **-1.48 ± 0.03** |
| | Run 2 | 50 ns | 17.41 ± 0.05 | 19.49 ± 0.07 | |
| | Run 3 | 50 ns | 17.36 ± 0.04 | 18.03 ± 0.04 | |
| | **Average** | | **17.40 ± 0.02** | **18.88 ± 0.03** | |
| **S766A** | Run 1 | 50 ns | -4.38 ± 0.04 | -4.55 ± 0.06 | **0.16 ± 0.04** |
| | Run 2 | 50 ns | -4.38 ± 0.05 | -4.52 ± 0.05 | |
| | Run 3 | 50 ns | -4.44 ± 0.04 | -4.62 ± 0.05 | |
| | **Average** | | **-4.40 ± 0.03** | **-4.56 ± 0.03** | |
| **A767S** | Run 1 | 50 ns | 4.64 ± 0.05 | 3.99±0.05 | **0.38 ± 0.04** |
| | Run 2 | 50 ns | 4.08 ± 0.05 | 3.79±0.04 | |
| | Run 3 | 50 ns | 4.28 ± 0.05 | 4.07±0.06 | |
| | **Average** | | **4.33 ± 0.03** | **3.95 ± 0.03** | |
| **I778A** | Run 1 | 50 ns | -5.53 ± 0.07 | -4.25 ± 0.06 | **-1.12 ± 0.05** |
| | Run 2 | 50 ns | -5.74 ± 0.06 | -4.66 ± 0.05 | |
| | Run 3 | 50 ns | -5.76 ± 0.05 | -4.75 ± 0.06 | |
| | **Average** | | **-5.67 ± 0.04** | **-4.55 ± 0.03** | |
| **L781A** | Run 1 | 50 ns | 14.13 ± 0.03 | 12.34 ± 0.03 | |
| | Run 2 | 50 ns | 14.15 ± 0.04 | 12.48 ± 0.04 | **1.76 ± 0.02** |
| | Run 3 | 50 ns | 14.07 ± 0.05 | 12.23 ± 0.03 | |
| | **Average** | | **14.11 ± 0.02** | **12.35 ± 0.02** | |

**Table S4:** Data similar to Table S3 are shown for single mutations in PDE5 (corresponding to variations observed in PDE6 isoforms).

| Mutation | Run | Sim. length | ΔG Comp | ΔG Apo | ΔΔG (kcal/mol) |
|---|---|---|---|---|---|
| **F564L** | Run 1 | 50 ns | -19.13 ± 0.07 | -19.59 ± 0.06 | **0.50 ± 0.05** |
| | Run 2 | 50 ns | -19.31 ± 0.07 | -19.64 ± 0.06 | |
| | Run 3 | 50 ns | -18.89 ± 0.09 | -19.62 ± 0.06 | |
| | **Average** | | **-19.11 ± 0.04** | **-19.61 ± 0.03** | |
| **R616Q*** | Run 1 | 50 ns | 209.29 ± 0.18 | 207.72 ± 0.18 | **1.49 ± 0.14** |
| | Run 2 | 50 ns | 209.07 ± 0.20 | 207.10 ± 0.22 | |
| | Run 3 | 50 ns | 208.61 ± 0.18 | 207.68 ± 0.16 | |
| | **Average** | | **208.99 ± 0.10** | **207.50 ± 0.10** | |
| **I778V** | Run 1 | 50 ns | -5.22 ± 0.06 | -4.33 ± 0.04 | **-0.69 ± 0.03** |
| | Run 2 | 50 ns | -5.00 ± 0.08 | -4.58 ± 0.05 | |
| | Run 3 | 50 ns | -4.81 ± 0.05 | -4.06 ± 0.04 | |
| | **Average** | | **-5.01 ± 0.03** | **-4.32 ± 0.02** | |
| **L781M** | Run 1 | 50 ns | 10.85 ± 0.11 | 10.12 ± 0.13 | **0.41 ± 0.07** |
| | Run 2 | 50 ns | 10.49 ± 0.10 | 10.49 ± 0.10 | |
| | Run 3 | 50 ns | 11.01 ± 0.10 | 10.52 ± 0.12 | |
| | **Average** | | **10.78 ± 0.05** | **10.37 ± 0.06** | |

**Table S5:** Data similar to Table S3 are shown for double/triple mutations in PDE5 (corresponding to variations observed in PDE6 isoforms).

| Mutation | Run | Sim. length | ΔG Comp | ΔG Apo | ΔΔG (kcal/mol) |
|---|---|---|---|---|---|
| **I778V + L781M [PDE6 CONE]** | Run 1 | 50 ns | 6.65 ± 0.12 | 5.91 ± 0.10 | **0.71 ± 0.07** |
| | Run 2 | 50 ns | 6.90 ± 0.10 | 5.81 ± 0.11 | |
| | Run 3 | 50 ns | 6.00 ± 0.13 | 5.68 ± 0.10 | |
| | **Average** | | **6.51 ± 0.06** | **5.80 ± 0.05** | |
| **I778V + F564L [PDE6 ROD]** | Run 1 | 50 ns | -23.21 ± 0.10 | -24.7 ± 0.11 | **1.12 ± 0.07** |
| | Run 2 | 50 ns | -23.86 ± 0.14 | -24.43 ± 0.09 | |
| | Run 3 | 50 ns | -23.24 ± 0.08 | -24.53 ± 0.11 | |
| | **Average** | | **-23.43 ± 0.06** | **-24.55 ± 0.05** | |
| **I778V + L781M + R616Q* [PDE6 CONE]*** | Run 1 | 50 ns | 218.64 ± 0.22 | 215.62 ± 0.22 | **2.29 ± 0.18** |
| | Run 2 | 50 ns | 218.11 ± 0.29 | 215.12 ± 0.18 | |
| | Run 3 | 50 ns | 216.14 ± 0.27 | 215.28 ± 0.22 | |
| | **Average** | | **217.63 ± 0.15** | **215.34 ± 0.11** | |

*PDE6 variant used in the experimental study reporting evodiamine derivatives as PDE5 allosteric inhibitors.

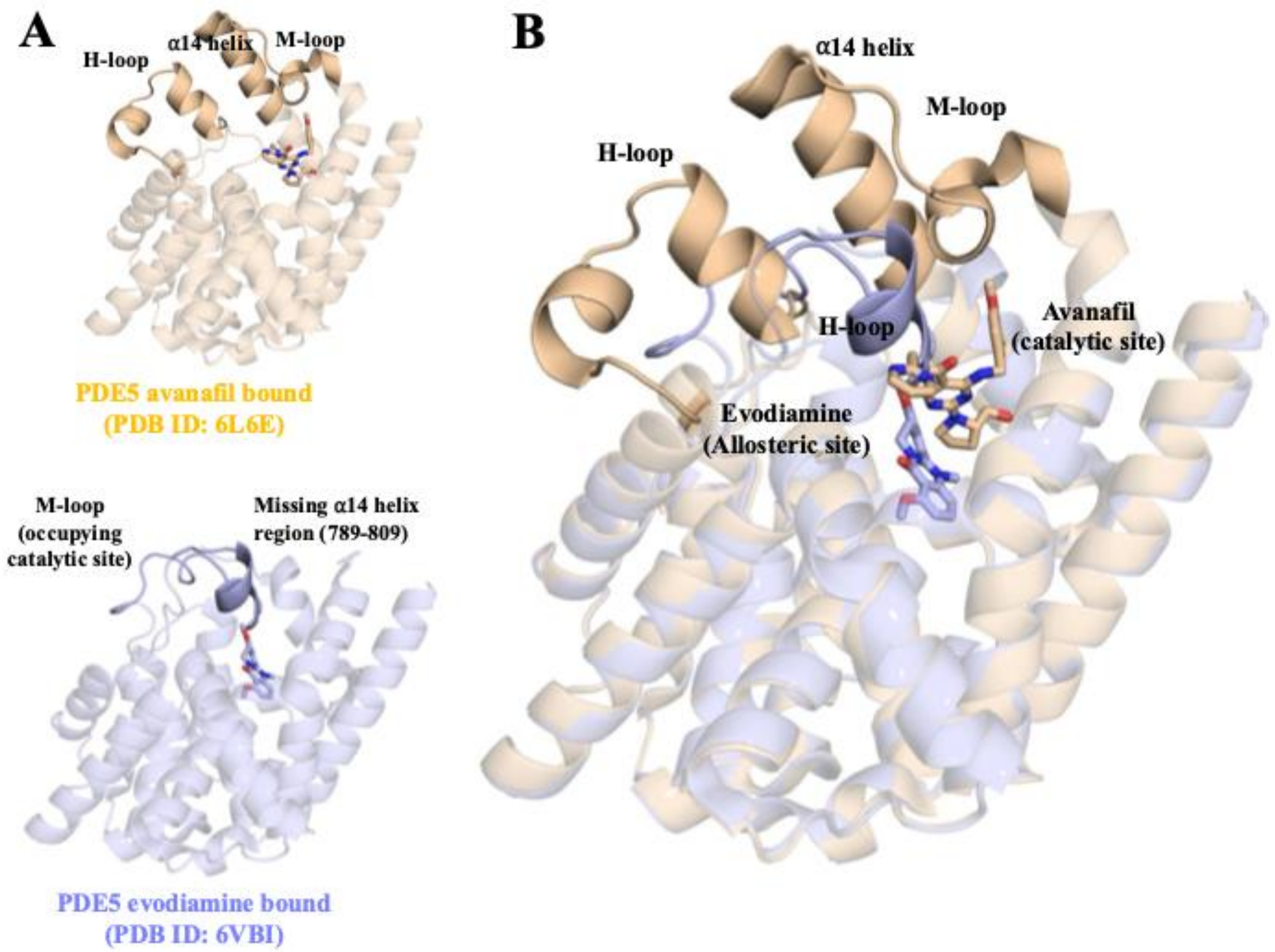


**Figure S1: Structural comparison of allosteric and orthosteric inhibitor binding sites in PDE5.** (A) Structural representations of PDE5 bound to the allosteric inhibitor evodiamine (EVO; PDB ID: 6VBI) and the orthosteric inhibitor avanafil (PDB ID: 6L6E). (B) Structural superposition of PDE5 and inhibitor structures. The PDE5 protein is shown in cartoon representation, highlighting key regulatory elements including the α14-helix, H-loop, and M-loop. EVO occupies a distinct allosteric pocket adjacent to the regulatory helices, whereas avanafil binds within the conserved catalytic site. The region corresponding to residues 789–809 of the α14-helix, which lacks electron density in the EVO-bound crystal structure, is indicated.

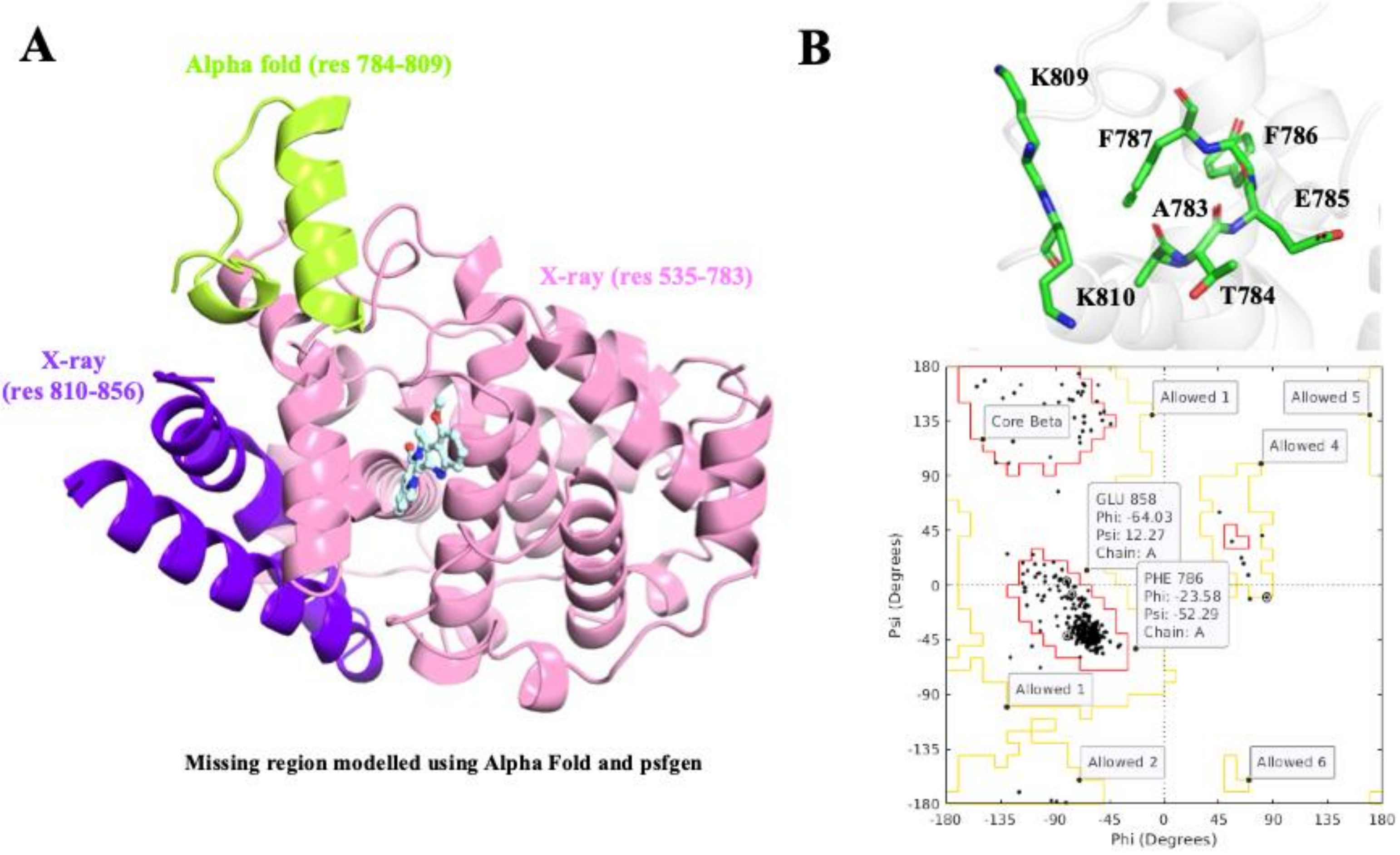


**Figure S2: Structural modeling of the missing α14-helix/M-loop region in the PDE5–EVO complex.** (A) Structures used in Integrative modeling to reconstruct residues 784–809, which are absent in the PDE5 crystal structure (PDB ID: 6VBI). The missing segment was extracted from the AlphaFold-predicted full-length PDE5 model and assembled between the resolved X-ray fragments (residues 535–783 and 810–856) using psfgen. (B) Structural validation of the reconstructed model showing connectivity between the crystallographic and AlphaFold-derived regions. Ramachandran plot analysis confirmed that all residues in the modeled segment occupy favored or allowed regions, with no outliers.

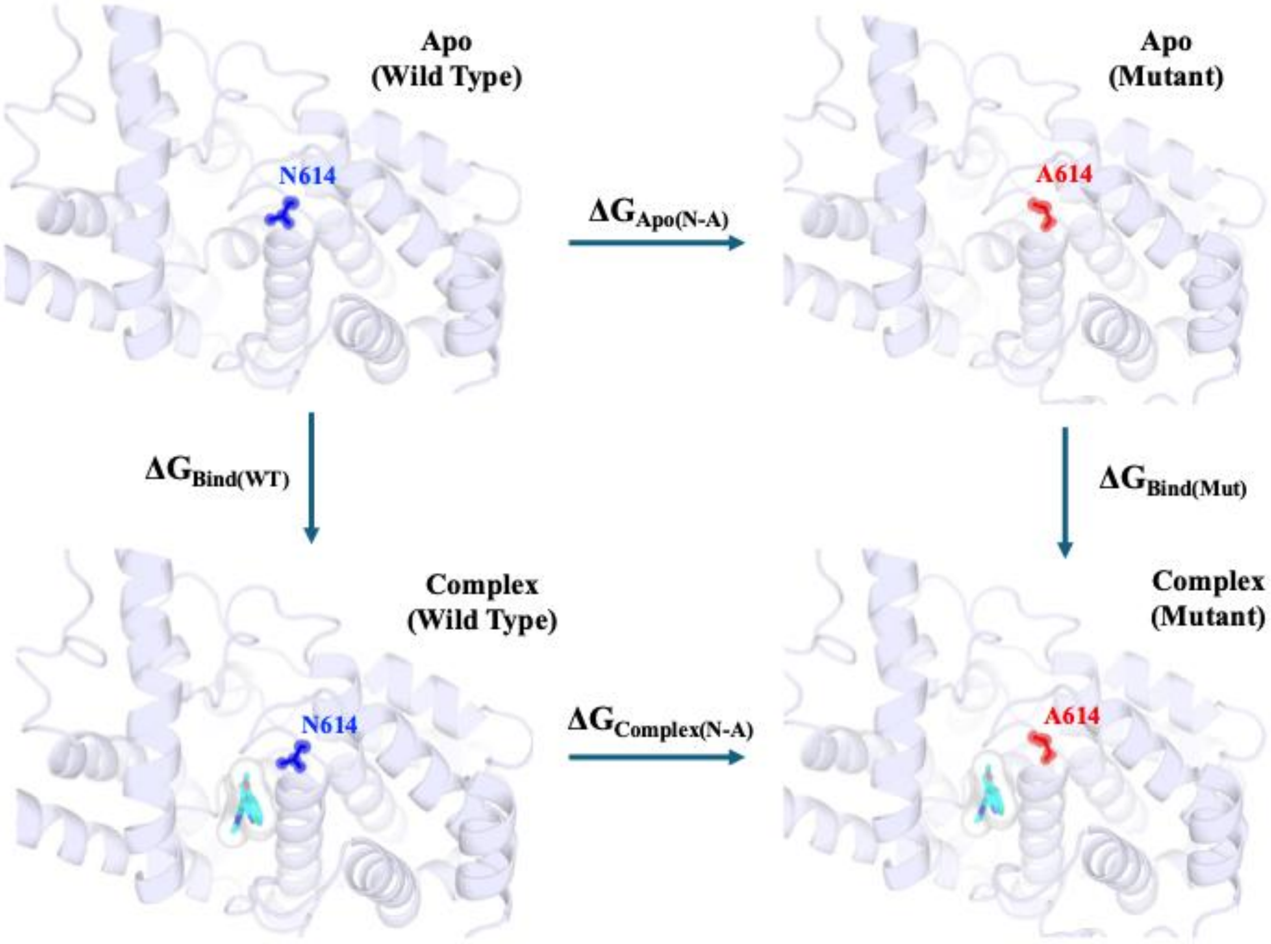


**Figure S3: The thermodynamic cycle for estimating the relative binding free energy (ΔΔG) of perturbations in PDE5.** The vertical arms of the thermodynamic cycle correspond to binding of the allosteric inhibitor (EVO) to PDE5, and the horizontal arms correspond to the alchemical transformation of a WT amino-acid into mutated one in complex (lower horizontal arm) when bound to the inhibitor and when free (upper horizontal arm) in water. The free energy perturbation method was used to compute the free energy changes (ΔG comp and ΔG Apo).

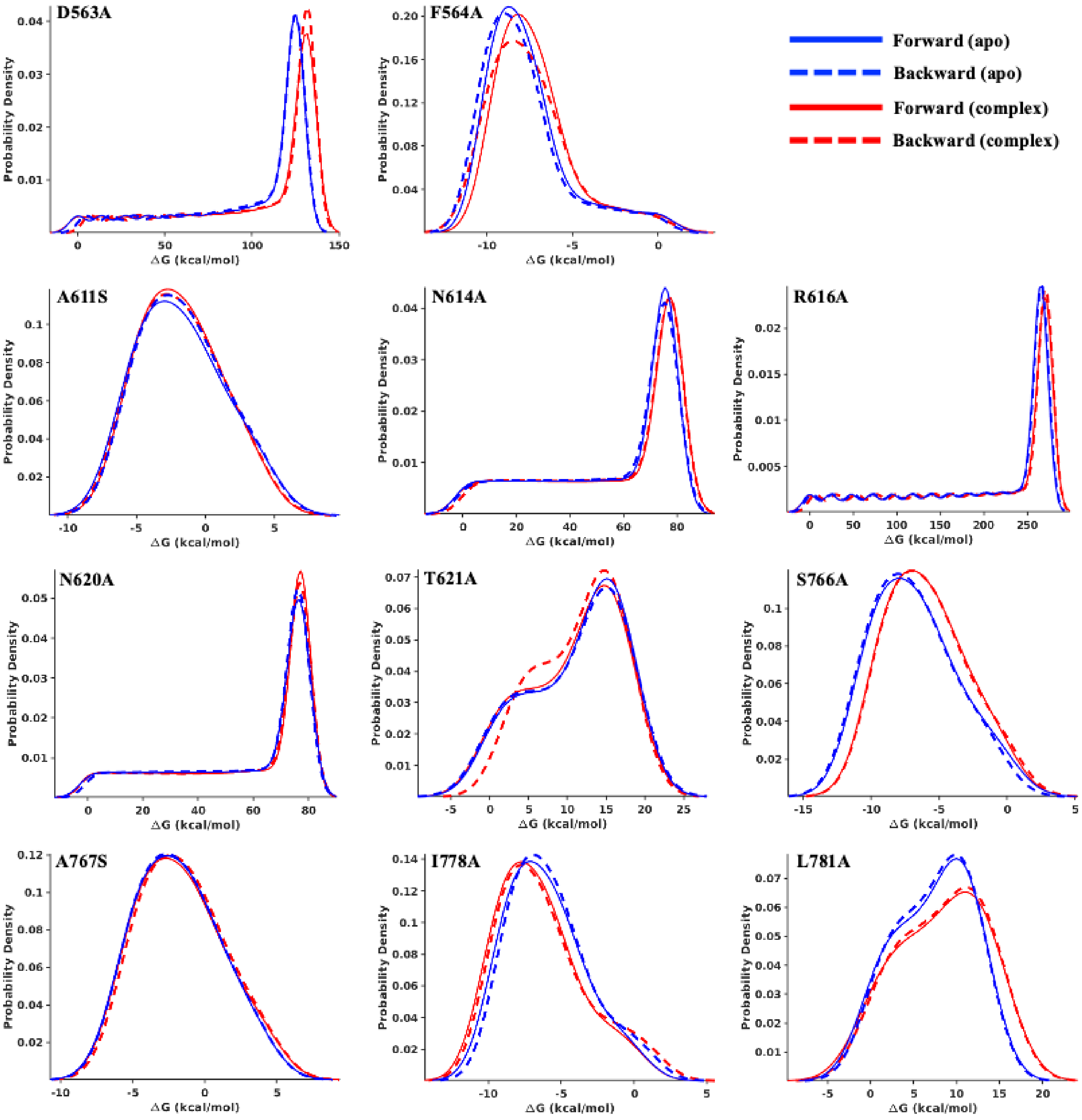


**Figure S4: Distributions of free energy (ΔG) for alanine mutations in PDE5–EVO complex.** The plots show the distributions of computed free energy (**ΔG**) for each transformation across all λ windows, for both the apo protein and inhibitor-bound complex. Forward and backward alchemical transformations are shown for each system. Data represent the cumulative results from three independent simulation replicas, with three forward and three backward transformations.

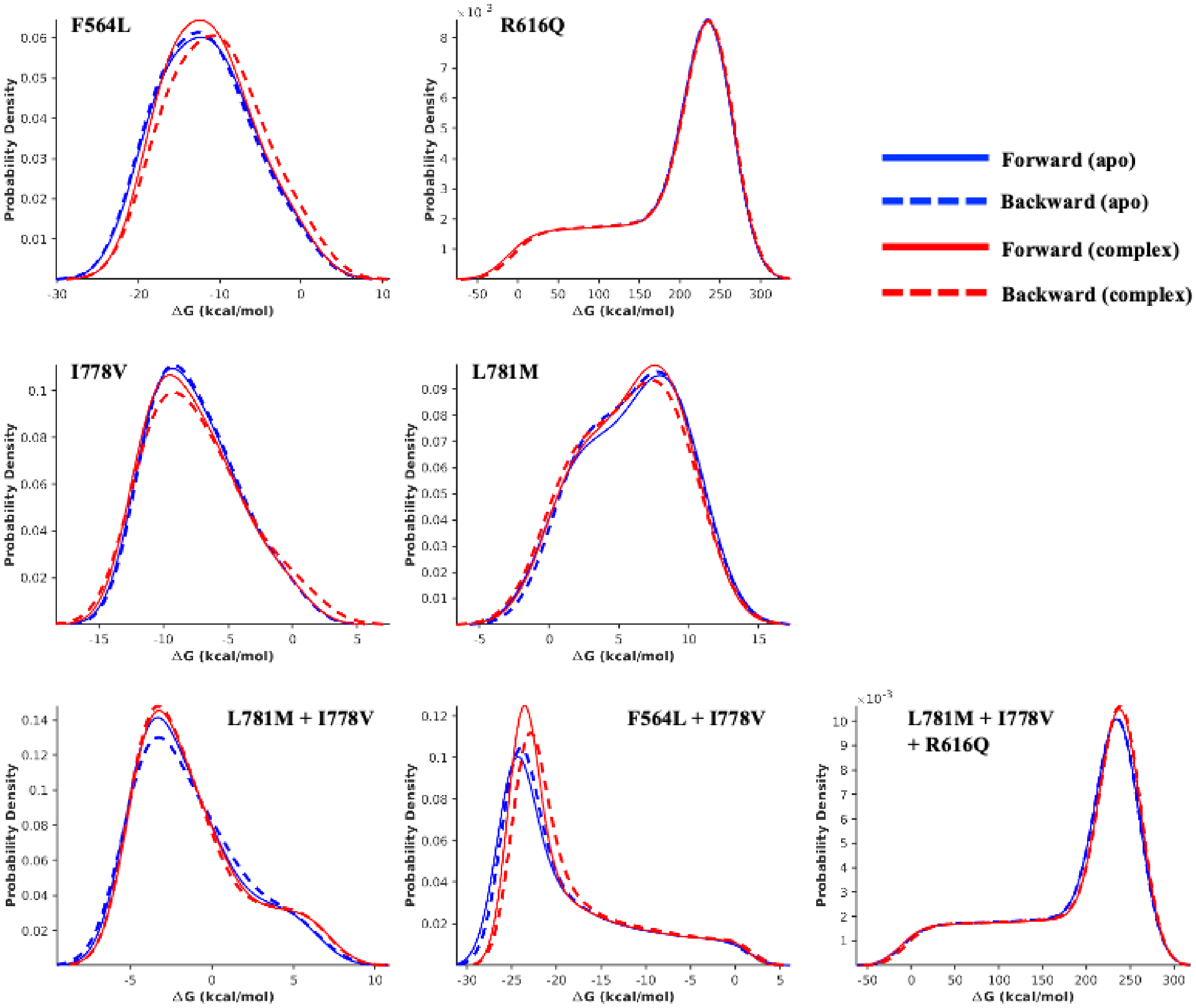


**Figure S5:** Data similar to Figure S4 are shown for single, double and triple mutations.

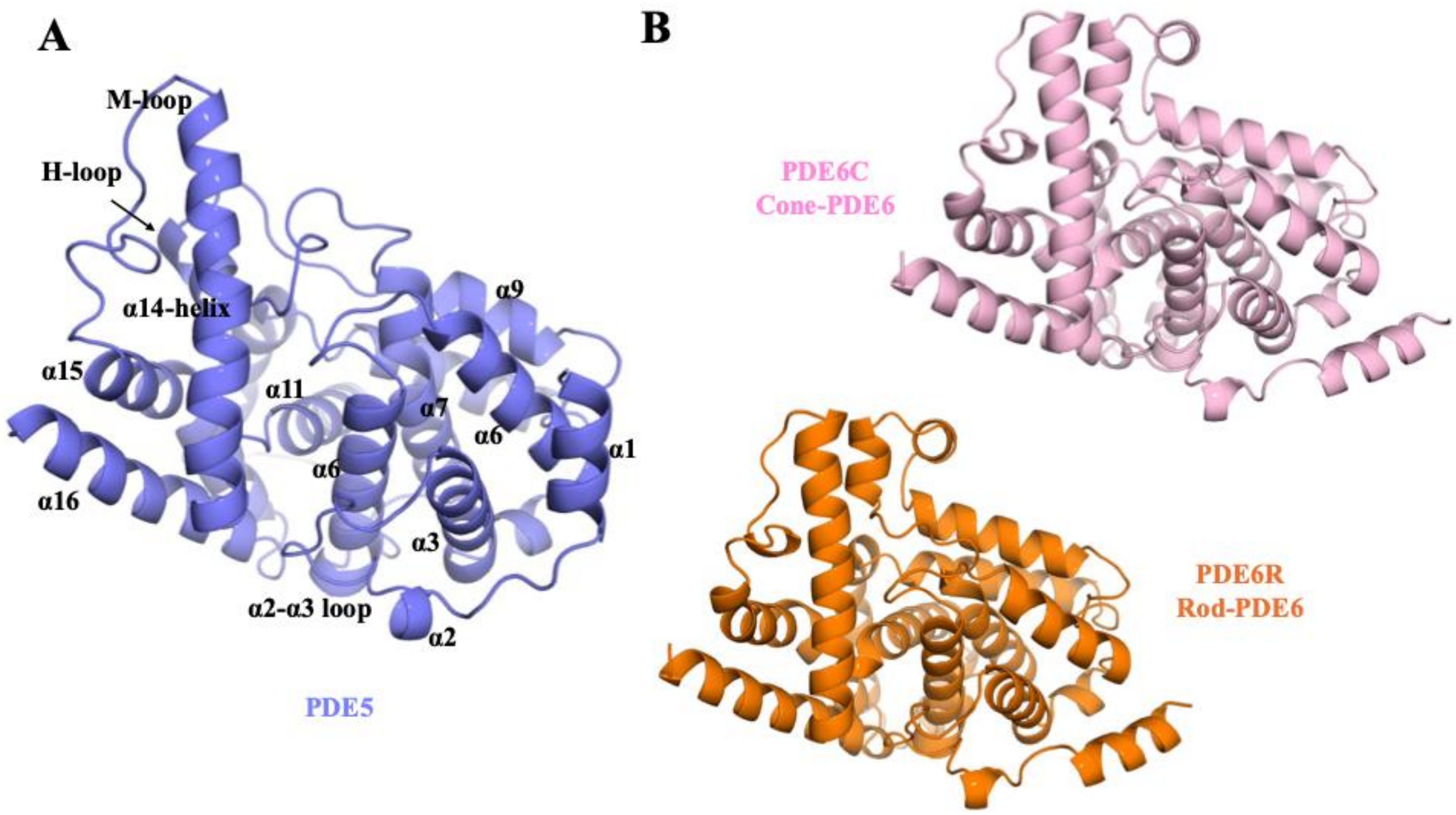


**Figure S6: Structural models of the catalytic domains of PDE5 and PDE6 isoforms.** (A) Modeled structure of PDE5 shown in the cartoon representation; the allosteric inhibitor EVO is omitted from the pocket. (B) Structural models of PDE6 isoforms (PDE6C; cone and PDE6R; rod). The PDE6 structures were predicted using AlphaFold3. The sequence information was obtained from UniProt entries P51160 (PDE6 cone) and P16499 (PDE6 rod).

*Note: The gamma (γ) subunit (integral part of PDE6 catalytic domain) was not modeled in this study. Only the alpha (α) subunit of the catalytic domain of PDE6 (both isoforms) was modeled to compare the allosteric binding pocket residues and perform structural alignment.*

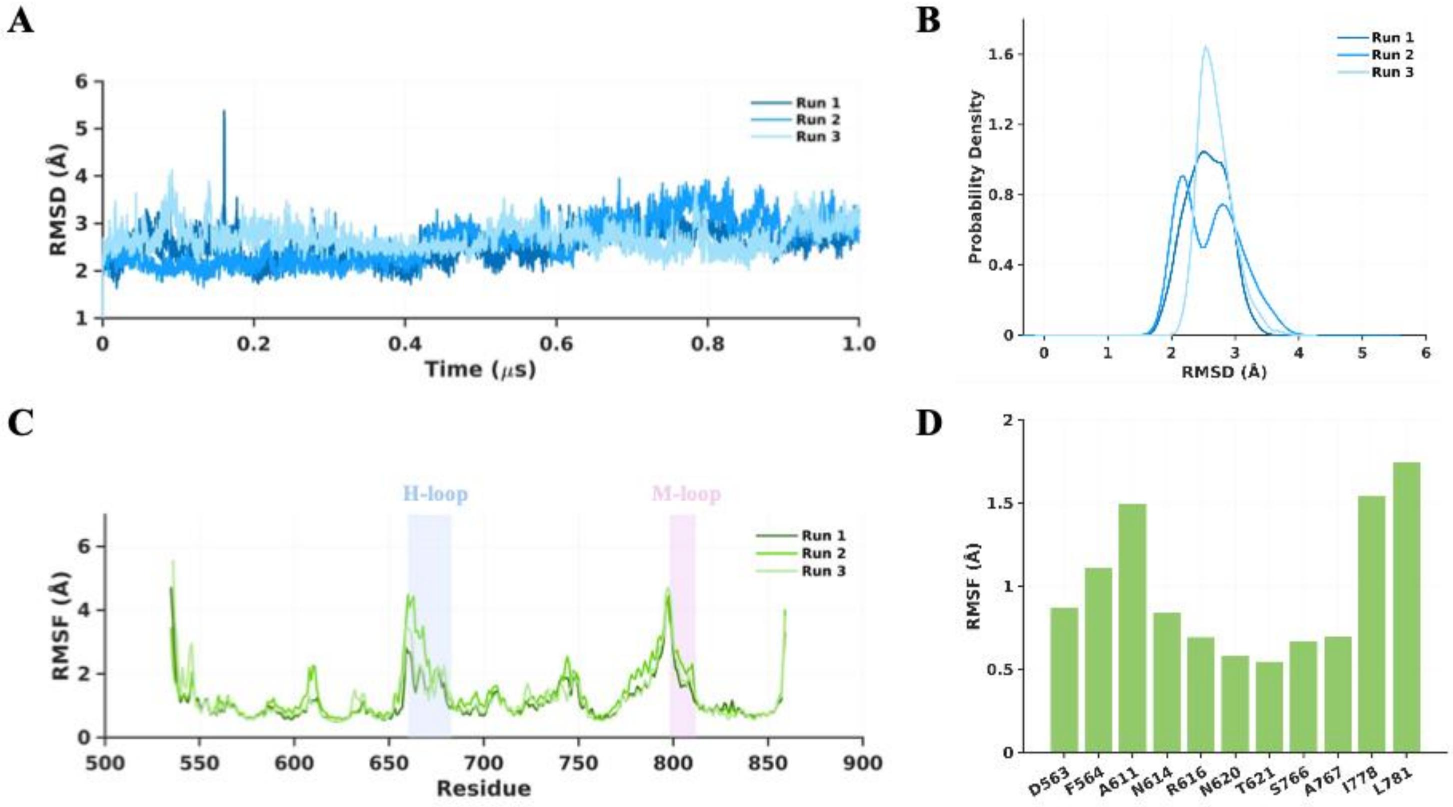


**Figure S7: Structural metrics root mean squared deviation (RMSD) and root mean squared fluctuation (RMSF) obtained from all atom MD simulations of the PDE5-EVO complex.** (A) RMSD of the protein backbone atoms. (B) Density distribution of RMSD of protein atoms. (C) RMSF of the $C_\alpha$ atoms. (D) RMSF for the allosteric pocket residues surrounding EVO in PDE5.

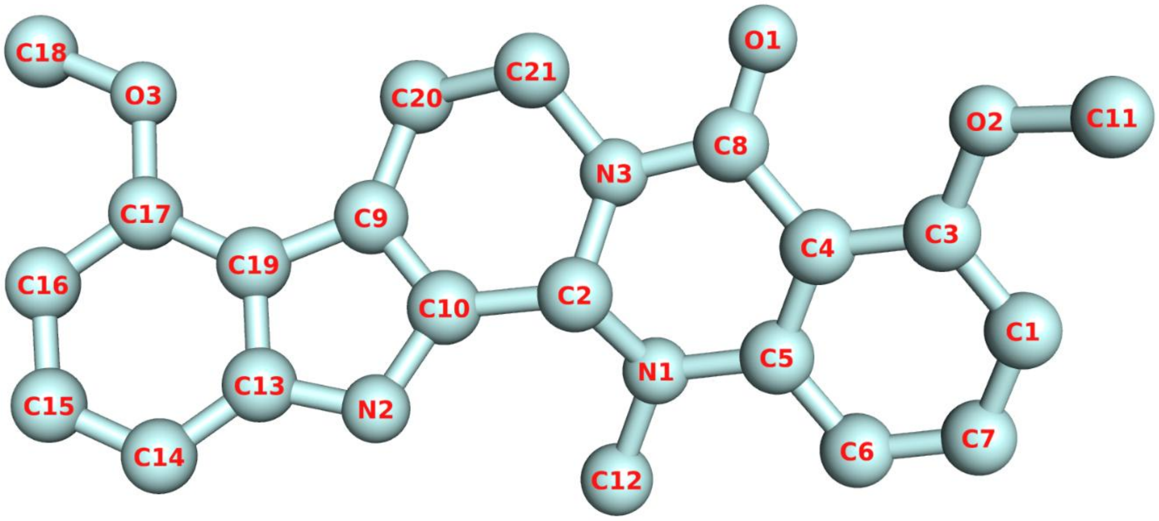


**Figure S8: Chemical structure of the Evodiamine derivative (EVO) highlighting the heavy atoms (C, N and O).** The atom names are labeled in red.
